# M18BP1 valency and a distributed interaction footprint determine epigenetic centromere specification in humans

**DOI:** 10.1101/2025.06.27.661933

**Authors:** Kai Walstein, Louisa Hill, Doro Vogt, Ingrid R. Vetter, Dongqing Pan, Andrea Musacchio

**Affiliations:** Department of Mechanistic Cell Biology, Max Planck Institute of Molecular Physiology, Otto-Hahn-Straße 11, 44227 Dortmund, Germany; Centre for Medical Biotechnology, Faculty of Biology, University Duisburg-Essen, Essen, Germany

**Keywords:** centromere, CENP-A, M18BP1, HJURP, MIS18, CCAN, kinetochore, CDK1

## Abstract

Deposition of CENP-A, the histone H3 variant considered an epigenetic landmark of centromeres, reflects cell-cycle-regulated assembly of M18BP1, HJURP, and PLK1 on a divalent MIS18α/β scaffold. The localization determinants of this machinery remain poorly characterized. Here we report that artificial M18BP1 dimerization bypasses MIS18α/β, allowing the identification of at least four determinants of M18BP1 centromere localization. These include the SANTA domain, of which we report the first structure, as well as linear motifs in disordered neighboring regions, of which we characterize the interaction footprint on the CENP-A associated 16-subunit constitutive centromere-associated network (CCAN). Our observations imply that M18BP1, after dimerization, is necessary and sufficient for centromere localization. Its cell-cycle-dependent dimerization on MIS18α/β promotes initial recognition of a multivalent centromeric assembly of old CENP-A and associated proteins, followed by cooption of PLK1 and HJURP and new CENP-A deposition. Our results shed new light on the determinants of centromere epigenetic inheritance in humans.

## Introduction

Chromosomes are carriers of the genome, and their faithful distribution to the daughter cells during a mother cell’s equational or reductional divisions (mitosis and meiosis, respectively) is essential for the propagation of life. These processes begin with the assembly of a cytoskeletal structure in the mother cell known as the spindle. Chromosomes interact with spindle microtubules through kinetochores, macromolecular complexes built on dedicated chromosome loci known as centromeres (Musacchio and Desai, 2017). Attainment of chromosome biorientation licenses mitotic exit, allowing the irreversible separation of chromosomes and their segregation into the daughter cells. The histone H3 variant centromere protein A (CENP-A) is an almost ubiquitous landmark of centromeres in eukaryotes (McKinley and Cheeseman, 2016; Mitra et al., 2020b). To build the kinetochore, CENP-A interacts with the constitutive centromere-associated network (CCAN), a supramolecular assembly of 16 proteins in human that is also largely conserved in evolution. The CCAN, in turn, recruits the components of the outer kinetochore, which form the microtubule-binding interface (McKinley and Cheeseman, 2016; Musacchio and Desai, 2017).

What ensures that centromeres maintain their size and position on chromosomes is a crucial and only partly answered question. Because CENP-A undergoes a 2-fold dilution during DNA replication, when it is redistributed to the sister chromatids, the reduction of its levels must be compensated through new CENP-A deposition. Foundational work demonstrated that this occurs in the G1 phase of the cell cycle, and thus before the dilution of CENP-A during S-phase (Jansen et al., 2007; Schuh et al., 2007). In primates, including humans, CENP-A associates with an abundant 171-bp DNA repeat known as α-satellite, which was therefore initially identified as a putative genetic determinant of centromeres (McKinley and Cheeseman, 2016; Mitra *et al*., 2020b). As new CENP-A is deposited near the old CENP-A, it might be expected to be directly attracted to vacant α-satellite repeats. However, the great variability of repeat sequences in different organism, and the discovery in several organisms of chromosomes with centromeres established on non-repetitive but diverse DNA sequences, forced to abandon the idea of a strict relationship between DNA sequence and centromere identity, in favor of a model in which a self-sustaining protein-based template maintains centromere identity through subsequent cell divisions (McKinley and Cheeseman, 2016; Mitra *et al*., 2020b). Reconstitution of ectopic centromeres after forcing repositioning of CENP-A or other CCAN subunits proved an important prediction of this model (Barnhart et al., 2011; Gascoigne et al., 2011; Hori et al., 2013; Mendiburo et al., 2011).

Further support to an epigenetic model of centromere specification based on a self-sustaining protein-based template came from the discovery of dedicated CENP-A deposition machinery (Mitra *et al*., 2020b; Stellfox et al., 2013). In humans, this machinery includes the MIS18 complex (a 4:2 hexamer of the MIS18α and MIS18β subunits, henceforth MIS18α/β), M18BP1 (also known as KNL2), the specialized CENP-A:H4 chaperone HJURP, and Polo-like kinase 1 (PLK1) (Dunleavy et al., 2009; Foltz et al., 2009; Fujita et al., 2007; Hayashi et al., 2004; Maddox et al., 2007; McKinley and Cheeseman, 2014). Interaction of these proteins is disallowed by cyclin-dependent kinase (CDK) phosphorylation from late G1-phase to M-phase (Conti et al., 2024; McKinley and Cheeseman, 2014; Pan et al., 2017; Parashara et al., 2024; Silva et al., 2012; Spiller et al., 2017; Stankovic et al., 2017) (Figure 1A). Rapid decline of CDK activity at the metaphase-to-anaphase transition permits interactions among these proteins that promote their recruitment and activation at centromeres and the discharge of new CENP-A near the existing CENP-A pool (Jansen *et al*., 2007). Thus, CENP-A deposition occurs during the G1 phase of the cell cycle, before DNA replication, contrary to the replenishment of H3:H4, which occurs concomitantly with its dilution during DNA replication (Alabert et al., 2015; Xu et al., 2010).

**Figure 1.**
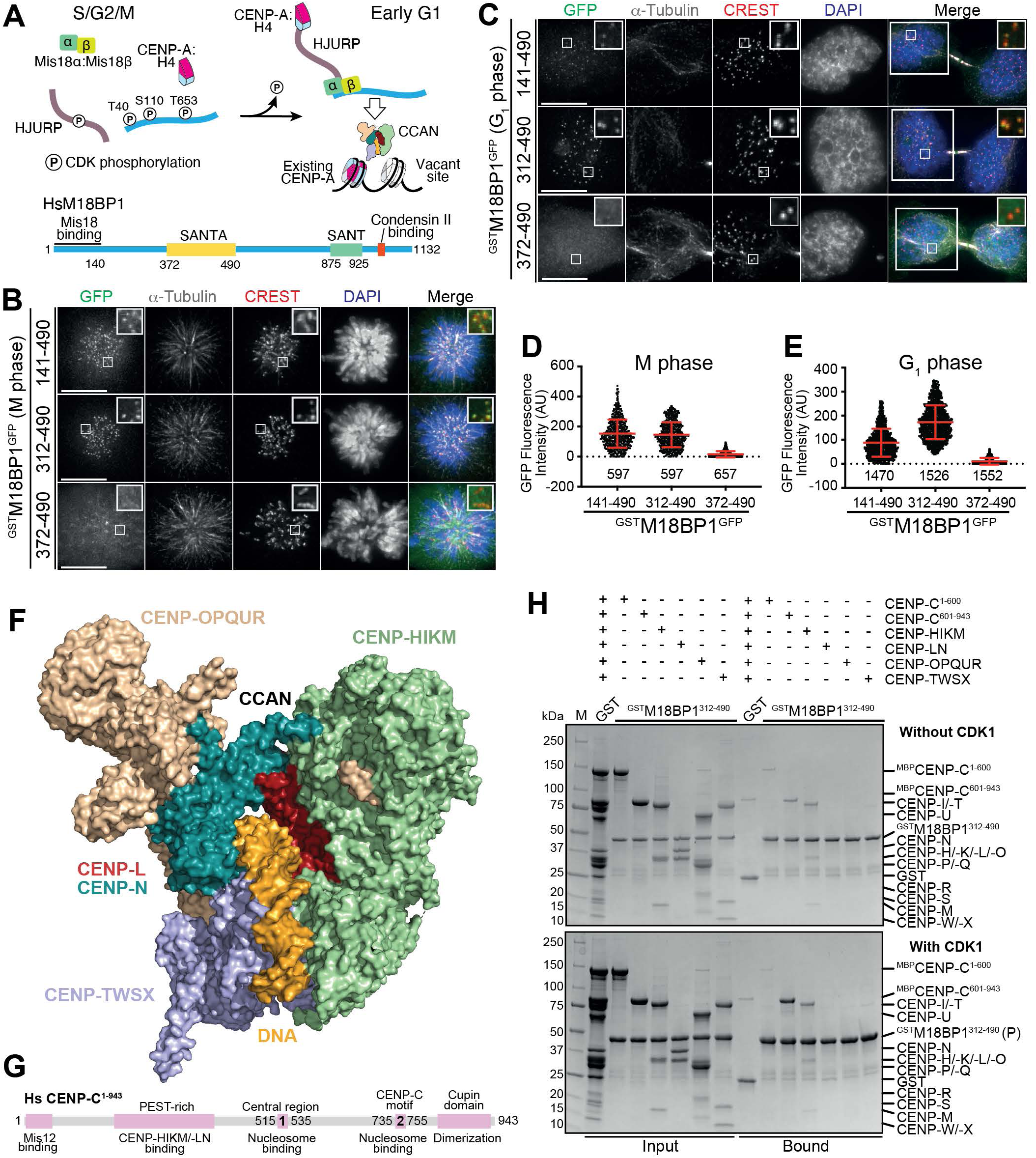
Identification of a minimal centromere targeting module of M18BP1. (**A**) Schematic model of the CDK phosphorylation-regulated assembly of the Mis18 complex and its centromeric recruitment in G1. HJURP is phosphorylated at multiple sites during mitosis (Flores Servin *et al*., 2023; Muller et al., 2014; Stankovic *et al*., 2017; Wang et al., 2014). (**B**) and (**C**) Representative images of fixed HeLa cells showing GFP fluorescence of indicated stably expressed M18BP1 variants and immuno-stained α-tubulin in mitosis and G1. Centromeres were visualized by CREST sera, and DNA was stained by DAPI. White scale bars indicate 10 µm. (**D**) and (**E**) Quantification of the centromeric GFP-fluorescence intensities of the indicated expressed M18BP1 variants in mitosis and G1. Centromeres were detected in the CREST channel using a script for semiautomated quantification. Red bars indicate mean and standard deviation of all quantified centromere fluorescence intensities from three independent experiments. Here and in other equivalent panels reporting quantifications of fluorescence intensity, the number of counted kinetochores is indicated. (**F**) Structural model of the constitutive centromere associated network (CCAN) bound to DNA (PDB TR5S). (**G**) Organization of human CENP-C. The arrangement of the binding sites recapitulates the outer-to-inner kinetochore axis. (**H**) GSH-resin pull-down assay with GST (negative control) and ^GST^M18BP1^312-490^ as baits. The added CCAN subcomplexes are indicated above each lane. The experiment was performed in the absence (upper gel) and in the presence (lower gel) of CCC kinase complex.

The epigenetic specification model puts emphasis on the protein-based interactions between centromeres and CENP-A loading machinery (herewith CALM) that promotes CENP-A deposition at the appropriate time and place. Nonetheless, the precise features that allow the CALM to become recruited specifically to the centromere remain largely elusive. In previous work, specific subunits of the CCAN, most notably CENP-C and CENP-I, were implicated in centromere recruitment of the CALM (Dambacher et al., 2012; French and Straight, 2019; Hoffmann et al., 2020; McKinley and Cheeseman, 2014; Mitra et al., 2020a; Moree et al., 2011; Shono et al., 2015). However, both subunits are also important for CCAN stability (Basilico et al., 2014; Guo et al., 2017; Klare et al., 2015; McKinley et al., 2015; Pesenti et al., 2022; Tachiwana et al., 2015; Walstein et al., 2021), and whether they are directly involved as CALM receptors or rather in the stabilization or recruitment of other CCAN subunits acting as CALM receptors remains unclear. On the CALM side, MIS18α/β has been proposed to cooperate with M18BP1 for kinetochore recruitment (Fujita *et al*., 2007; Pan *et al*., 2017; Stellfox et al., 2016). Furthermore, also HJURP was proposed to contain autonomous centromere-binding regions (Flores Servin et al., 2023; French et al., 2017; Tachiwana *et al*., 2015). Recent work, however, demonstrated that centromere recruitment of HJURP in early G1, at least in humans, is entirely contingent on its ability to interact with MIS18α/β under the control of PLK1 kinase activity (Conti *et al*., 2024; Parashara *et al*., 2024). Thus, collectively, the view of how the CALM is recruited to centromeres remains fragmented. Here, we shed light on this crucial question by mobilizing a combination of complex biochemical reconstitutions, cell biology in human cells, experimental structural work, and structural modelling.

## Results

### Sequence requirements for kinetochore recruitment of M18BP1

Residues 1-140 of human (Hs) M18BP1 (M18BP1^1-140^) are necessary for kinetochore recruitment of M18BP1 (Pan *et al*., 2017; Stellfox *et al*., 2016; Thamkachy et al., 2024). Two CDK phosphorylation sites within this fragment, T40 and S110, prevent binding to MIS18α/β. Their dephosphorylation upon mitotic exit allows binding of M18BP1 to MIS18α/β and robust centromere localization of the resulting octameric complex (McKinley and Cheeseman, 2014; Ohzeki et al., 2016; Pan *et al*., 2017; Silva *et al*., 2012; Spiller *et al*., 2017; Stankovic *et al*., 2017; Stellfox *et al*., 2016; Thamkachy *et al*., 2024). These observations are consistent with the idea that the centromere-binding determinants of M18BP1 are insufficient for robust centromere localization and need to be combined with those on MIS18α/β for stable recruitment (Stellfox *et al*., 2016). Indeed, M18BP1^141-1132^, which lacks the MIS18α/β binding region in M18BP1^1-140^, is not recruited to centromeres (Pan *et al*., 2017).

Nonetheless, we reasoned that an alternative interpretation of these observations is that dimerization of M18BP1 on MIS18α/β effectively makes it divalent, potentially increasing its kinetochore binding affinity (Erlendsson and Teilum, 2020; Mammen et al., 1998). This alternative interpretation would explain our puzzling previous observation that fusing M18BP1^141-1132^ to an N-terminal GST, a strong dimer, allowed robust kinetochore localization of the resulting construct despite its inability to bind MIS18α/β (Pan *et al*., 2017). In this view, phosphorylation of Thr40 and Ser110 may effectively control the valency of M18BP1 and its effective binding affinity for centromeres in G1.

To address this possibility, we asked if ectopic, cell-cycle invariant dimerization of M18BP1 through GST would allow us to identify localization determinants of M18BP1 and their cell cycle regulation. Towards this goal, we initially generated stable HeLa cell lines for inducible expression of various M18BP1 deletion mutants tagged with EGFP (and other tags as indicated). Constructs encompassing residues 1-490 of M18BP1 (M18BP1^1-490^) localized robustly to kinetochores in G1 phase, in agreement with a previous study (McKinley and Cheeseman, 2014), and regardless of artificial GST dimerization (Figure S1A-B, D). However, the GST-fused construct localized robustly also in mitosis, when the interaction with the MIS18 complex is suppressed by CDK phosphorylation (Figure S1C,E). This construct was expressed at much lower levels relative to its counterpart lacking GST (Figure S1A). Likely due to overexpression, the latter showed strong diffuse localization in addition to an unfocused kinetochore signal, in line with a previous study (McKinley and Cheeseman, 2014) (Figure S1C,E). Thus, binding determinants for the loading machinery at the kinetochore may exist not only in G1, but also in mitosis, suggesting that they are not cell-cycle regulated, contrary to the CENP-A loading machinery.

When gauged against our previous observation that the deleterious effects on kinetochore localization resulting from deleting residues 1-140 of M18BP1 are rescued by GST fusion, these results strongly suggest that residues 141-490 of M18BP1 contain crucial kinetochore binding determinants. To further characterize these determinants, we built three additional stable cell lines expressing segments 141-490, 312-490, and 372-490 fused to GST and EGFP (Figure S1F) and assessed their localization in G1- and M-phase. ^GST^M18BP1^141-490-EGFP^ and ^GST^M18BP1^312-490-EGFP^, but not ^GST^M18BP1^372-490-EGFP^, localized robustly to mitotic kinetochores (Figure 1B, D). Similarly, ^GST^M18BP1^141-490-EGFP^ and ^GST^M18BP1^312-490-EGFP^, but not ^GST^M18BP1^372-490-EGFP^, localized to kinetochores in G1, with ^GST^M18BP1^312-490-EGFP^ showing even more robust localization (Figure 1C, E). Thus, residues 312-490 of M18BP1, which encompass the SANTA (SANT-associated) domain (372-490) and a segment N-terminal to the SANTA (pre-SANTA) predicted to lack structural order (312-371), are a minimal kinetochore-targeting module of M18BP1 when fused to GST.

### M18BP1^312-490^ binds the CCAN

We harnessed *in vitro* biochemistry to identify the binding determinants of M18BP1^312-490^ within the kinetochore. In solid phase binding assays, M18BP1^312-490^ bound to CENP-A and H3 nucleosome core particles (H3^NCPs^ and CENP-A^NCPs^) indistinguishably (Figure S2A). Furthermore, incorporation of two centromere-specific histone marks, Histone H4 Lys20 monomethylation or α-amino-trimethylation of the CENP-A N-terminus (H4-K20me1 or CENP-A-Nme3, introduced enzymatically with SETD8 and NRMT1, respectively, in presence of S-adenosyl methionine) (Hori et al., 2014; Sathyan et al., 2017), did not affect the apparent affinity of M18BP1^312-490^ for CENP-A nucleosomes in electrophoretic mobility shift assays (EMSAs) (Figure S2B-D). In EMSA assays, M18BP1^312-490^ also bound DNA (Figure S2E). These interactions, whose likely primary determinant is unspecific DNA binding, are unlikely to explain the exquisitely specific kinetochore localization of M18BP1^312-490^.

We therefore turned onto the CCAN, which associates with CENP-A at the centromere-kinetochore interface. CCAN consists of several stable subcomplexes (Figure 1F). Its interaction with the CENP-A nucleosome is mediated by CENP-C, a 943-residue disordered protein with multiple interaction motifs disseminated along its length (Figure 1G). We asked whether recombinant CCAN subunits or subcomplexes interacted with GST-M18BP1^312-490^ immobilized on solid phase (Walstein *et al*., 2021; Weir et al., 2016). These assays identified interactions with the C-terminal region of CENP-C (CENP-C^601-943^, also indicated as CENP-C^C^) and with the CENP-HIKM complex. No binding or only weak binding was observed with CENP-C^1-600^, CENP-LN, CENP-OPQUR, or CENP-TWSX (Figure 1H, *upper panel*). Phosphorylation of the CCAN subunits with CDK1/Cyclin B/CKS1 kinase (CCC complex) and ATP (Huis In ’t Veld et al., 2022) caused a strong enhancement of the interaction with CENP-C^C^ (Figure 1H, lower panel), suggesting regulation of this interaction by mitotic phosphorylation (see below). Further dissection demonstrated that the entire M18BP1^312-490^ segment was required for robust binding to CENP-C^601-943^, as neither of two sub-fragments, M18BP1^312-371^ and M18BP1^372-490^ fused with GST and immobilized on solid phase bound it robustly (Figure 2A). Conversely, ^GST^M18BP1^312-371^ was sufficient to bind the CENP-HIKM complex, whereas no binding to ^GST^M18BP1^372-490^ (SANTA domain) was observed (Figure 2B).

**Figure 2.**
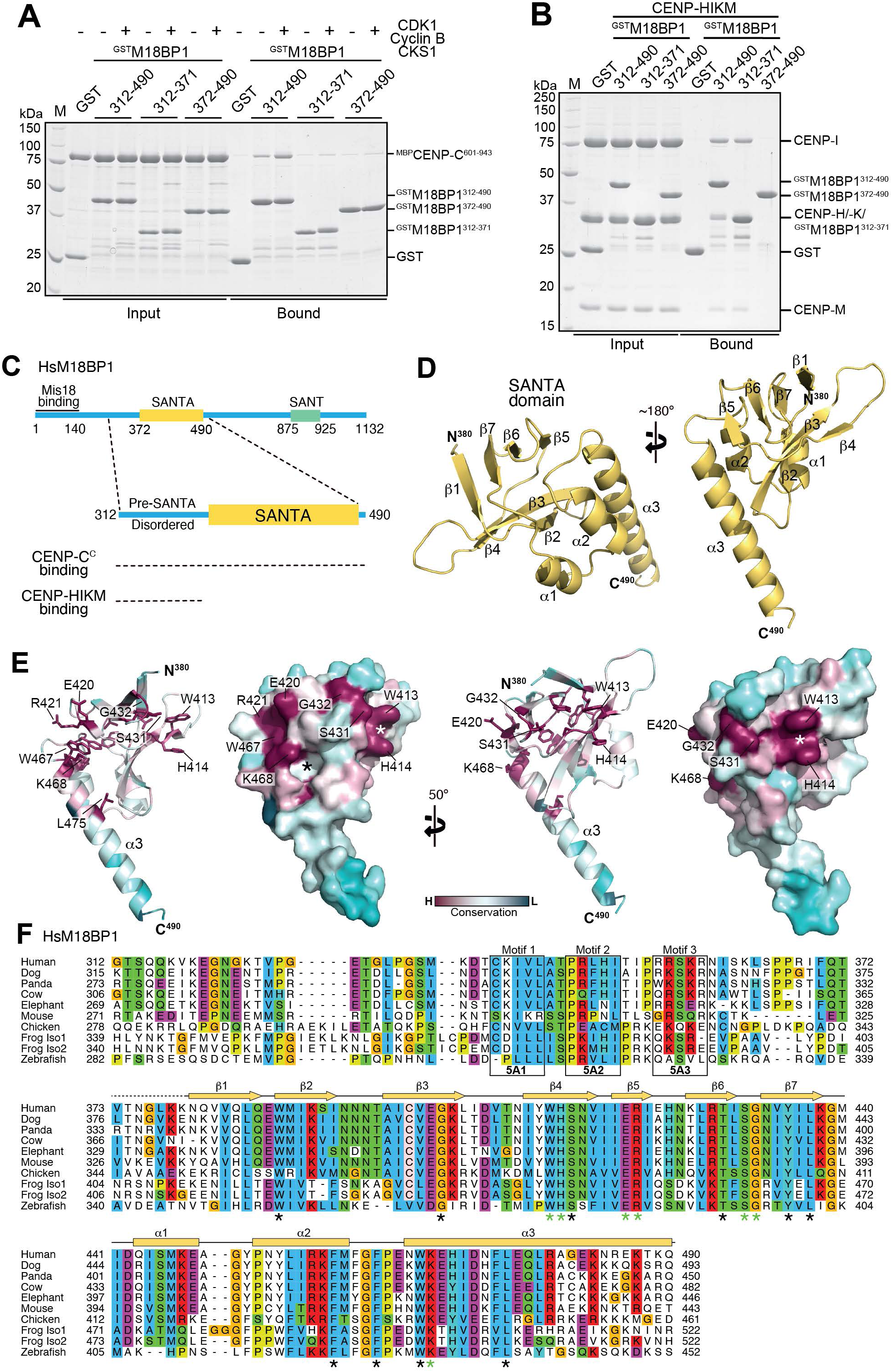
The SANTA domain and its N-Terminal extension provide distinct binding sites for the CCAN. (**A**) GSH-resin pull-down assay with GST (negative control) and ^GST^M18BP1 fragments as baits testing the binding to ^MBP^CENP-C^601-943^. The M18BP1 fragments used as baits and the addition of CCC kinase complex are indicated above each lane. (**B**) GSH-resin pull-down assay with GST (negative control) and ^GST^M18BP1 fragments as baits testing the binding to CENP-HIKM complex. The M18BP1 fragments used as baits are indicated above each lane. (**C**) Graphical summary of the obtained results in panels A and B. (D) Crystal structure of the human M18BP1 SANTA domain depicting the folded region from residue 380-490. The SANTA domain was crystallized as the fusion protein construct EGFP-**AS**-SANTA-**LEGT**-anti-GFP Nanobody-GGHHHHHH (see Methods). Only the SANTA moiety is shown, while the rest was omitted from the representation. (**E**) Structure of the SANTA domain. Conservation is indicated by a color code ranging from high (magenta) to low (turquoise) conservation from sequences in the alignment in panel (F). A prominent hydrophobic cavity and the exposed side chains of Trp413 and His414 are indicated with a black and a white asterisk, respectively. (**F**) Multiple sequence alignment showing conserved residues within M18BP1^312-490^ across mammalian and non-mammalian vertebrate species. Conserved residues in the hydrophobic core and on the surface are indicated with black and green asterisk, respectively.

These results implicate the disordered segment M18BP1^312-371^ (herewith also referred to as pre-SANTA region) in CENP-C^601-943^ and CENP-HIKM binding. They also implicate the SANTA domain in CENP-C^601-943^ binding, and indicate a potential role of mitotic CDK phosphorylation in modulating the interaction with CENP-C (summarized in Figure 2C).

### Crystal structure of the SANTA domain

We determined a crystal structure of a fusion construct of the SANTA domain (see Methods), the first for this domain class (Zhang et al., 2006), to a Bragg spacing of 2.4 Å (1 Å = 0.1 nm. See Table S1 for a summary of relevant crystallographic data). The folded region begins around residue 380 and ends at residue 490. It consists of a small globular domain formed by a pair of three- and four-stranded β-sheets packing against each other through a shared hydrophobic core. This core β-domain is followed by three consecutive α-helices, the third considerably longer than the first two, packing at one edge of the globule (Figure 2D). A search of the protein databank for structurally related domains DALI (Holm, 2022) did not identify folds closely related to the SANTA domain, while the Foldseek server (van Kempen et al., 2024) identified β-domains closely related to the SANTA’s in the predicted structures of several bacteriophages.

The SANTA domain displays several highly conserved residues, distributed both in its hydrophobic core or on its surface (Figure 2E. These residues are respectively indicated with black and green asterisks under the alignment in Figure 2F). Among the most prominent surface features are the parallel “tracks” of the exposed side chains of Trp413 and His414 on the β4 strand (indicated with a white asterisk in Figure 2E), as well as a prominent hydrophobic cavity (indicated with a black asterisk) delimited by the side chain of Lys468, at the beginning on the α3 helix. Together with the side chains of other highly conserved residues, including Glu420, Arg421, and Ser431, we predict these neighboring prominent conserved surface features to be part of a continuous, convex binding interface for an unknown target, likely an extended, flexible motif (see Discussion).

### Kinetochore-binding determinants of M18BP1^312-490^

To probe the significance of the interactions of M18BP1^312-490^ with CCAN for kinetochore recruitment of M18BP1, we introduced mutations in the pre-SANTA region and SANTA domain. Specifically, we created a double alanine mutant of Trp413 and His414 (the WH/AA mutant) to try impair the function of the SANTA domain. We also identified three short conserved motifs in the pre-SANTA region (Motifs 1-3) and generated three five-alanine mutants, named respectively 5A1, 5A2, and 5A3 (Figure 2F) to inactivate them. We introduced these mutations in the ^GST^M18BP1^312-490-EGFP^ construct and generated stable cell lines (Figure 3A and Figure S1G). The 5A1, 5A3, and WH/AA mutations all impaired kinetochore recruitment of ^GST^M18BP1^312-490-EGFP^ in early G1 cells (Figure 3B, D). Albeit expressed at lower levels (Figure S1G), the 5A2 mutant retained the ability to decorate kinetochores. In mitosis, neither the 5A1 nor the WH/AA mutant decorated kinetochores, while both 5A2 and 5A3 did (Figure 3B, D). Thus, Motif 3 is necessary for interphase recruitment of ^GST^M18BP1^312-490-EGFP^, but dispensable for its mitotic localization. As the SANTA domain is essential for mitotic as well as interphase localization of ^GST^M18BP1^312-490-EGFP^, we suspect that its centromere target persists during the cell cycle.

**Figure 3.**
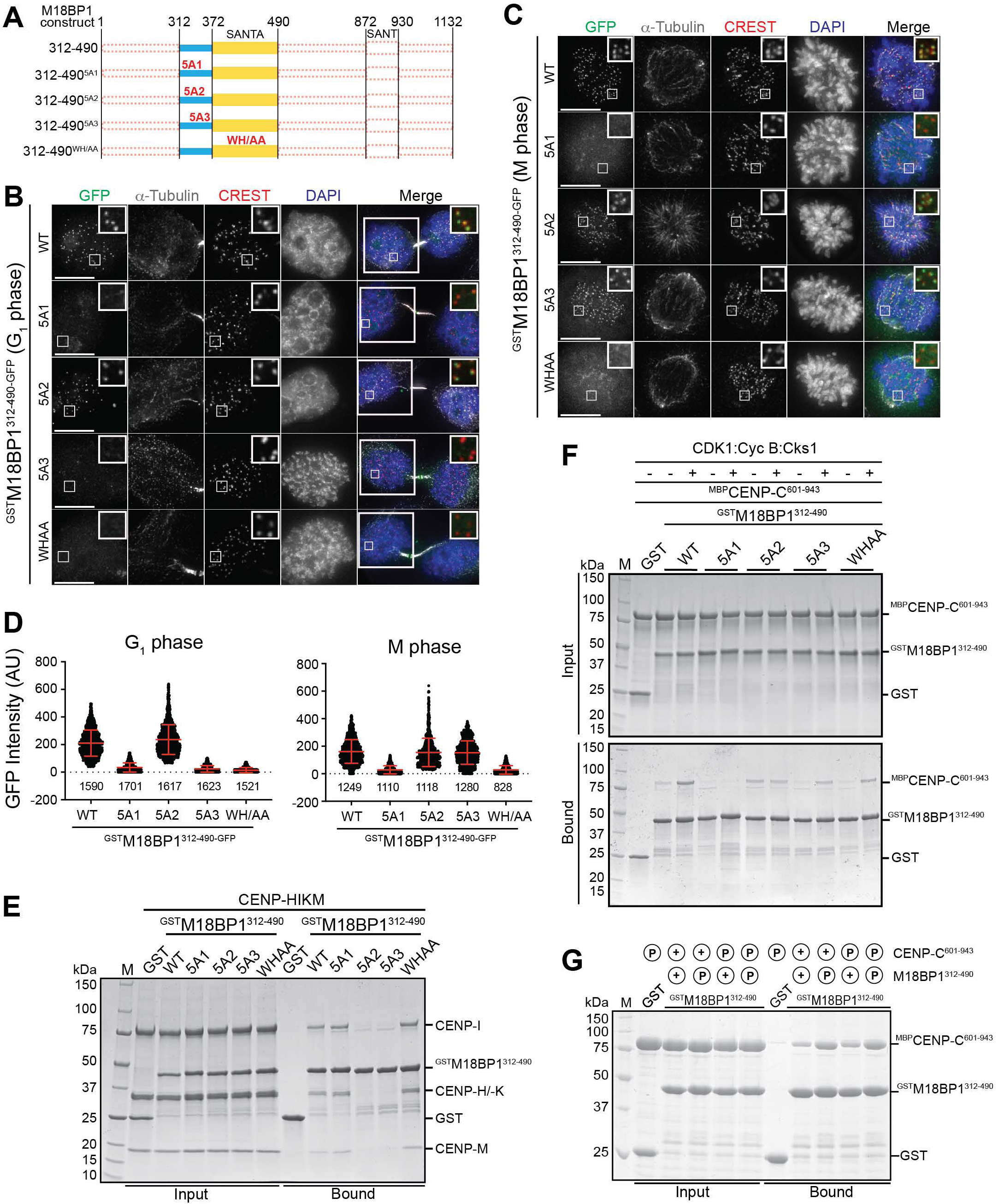
Unraveling the crucial interactions for mitotic and G1 localization of M18BP1. (**A**) Schematic showing the expressed ^GST^M18BP1^312-490-GFP^ variants (**B**) and (**C**) Representative images of fixed HeLa cells showing GFP fluorescence of indicated stably expressed M18BP1 variants and immuno-stained α-tubulin in G1 and mitosis, respectively. Centromeres were visualized by CREST sera, and DNA was stained by DAPI. White scale bars indicate 10 µm. (**D**) Quantification of the centromeric GFP-fluorescence intensities of the indicated expressed M18BP1 variants in G1 and mitosis, respectively. Centromeres were detected in the CREST channel using a script for semiautomated quantification. Red bars indicate mean and standard deviation of all quantified centromere fluorescence intensities from three independent experiments. (**E**) GSH-resin pull-down assay with GST (negative control) and ^GST^M18BP1^312-490^ variants as baits testing the binding to CENP-HIKM complex. The M18BP1 variants used as baits are indicated above each lane. (**F**) GSH-resin pull-down assay with GST (negative control) and ^GST^M18BP1^312-490^ variants as baits testing the binding to ^MBP^CENP-C^601-943^. The M18BP1 variants used as baits and the addition of CDK1:Cyclin B:Cks1 kinase complex are indicated above each lane (**G**) GSH-resin pull-down assay with GST (negative control) and ^GST^M18BP1^312-490^ as baits testing the binding to ^MBP^CENP-C^601-943^. In this experiment, ^GST^M18BP1^312-490^ and ^MBP^CENP-C^601-943^ were individually pre-phosphorylated by CDK1:Cyclin B:Cks1 kinase complex as indicated by a red encircled P above each lane.

We used *in vitro* binding assays to identify potential binding partners of Motifs 1-3 and of the SANTA domain. Initially, we immobilized wild type and mutant ^GST^M18BP1^312-490^ and tested their binding to the CENP-HIKM complex. Both 5A2 and 5A3 impaired binding of ^GST^M18BP1^312-490^ to CENP-HIKM, whereas 5A1 and the SANTA mutants bound normally (Figure 3E). To identify binding determinants of ^GST^M18BP1^312-490^ and CENP-HIKM, we performed crosslinking-mass spectrometry (XL-MS) experiments with CENP-HIKM and wild type ^GST^M18BP1^312-490^, using the 5A3 mutant as negative control (Figure S3A-B). ^GST^M18BP1^312-490^ crosslinked prominently with CENP-K, and less prominently with CENP-H and CENP-I, while the 5A3 mutant showed only few crosslinks. Albeit structurally stable, a CENP-HIKM mutant complex carrying substitutions at residues identified in the XL-MS experiment (CENP-I^Y488A^, CENP-H^WT^, CENP-K^S113R/G181R/E182K/E185K/D186R/E214K/E217K^, CENP-M^WT^) did not interact with ^GST^M18BP1^312-490^ in in vitro binding assays (Figure S3C). AlphaFold (AF) (Abramson et al., 2024; Jumper et al., 2021) predicts that Motifs 1 and 3 of M18BP1 form short helices while Motif 2 adopts an extended conformation (Figure S3D-E). Motif 2 may poke its hydrophobic side chains into a cradle between the CENP-H and CENP-K helices, while the positively charged residues of Motif 3 are predicted to interact with an acidic patch comprising Glu182, Glu185, Asp186, Glu214, and Glu217 of CENP-K (Figure S3D-E), which were all targeted in our CENP-HIKM mutant construct. Thus, the crosslinking data provide a good explanation of the effects of mutations in Motifs 2 and 3.

Next, we immobilized wild type and mutant ^GST^M18BP1^312-490^ and tested binding to ^MBP^CENP-C^601-943^ (Figure 3F). The 5A1 mutant appeared to disrupt CENP-C binding regardless of its CDK phosphorylation status. The 5A2, 5A3, and WH/AA mutants, on the other hand, bound immobilized ^MBP^CENP-C^601-943^, but less strongly than the wild type protein, with 5A2 apparently eliminating stimulation of binding by CDK phosphorylation. To analyze how CDK phosphorylation increases CENP-C^C^ binding, we pre-phosphorylated ^GST^M18BP1^312-490^ or ^MBP^CENP-C^601-943^, and only then tested binding to the unphosphorylated partner (Figure 3G). These results demonstrated unequivocally that the phosphorylation modulating the interaction is on ^GST^M18BP1^312-490^, as the phosphorylated ^MBP^CENP-C^601-943^ bound ^GST^M18BP1^312-490^ to the same levels observed in the absence of phosphorylation. Conversely, binding to ^MBP^CENP-C^601-943^ was strongly enhanced by CDK phosphorylation of ^GST^M18BP1^312-490^, whether or not ^MBP^CENP-C^601-943^ had been pre-phosphorylated (Figure 3G). The 5A3 mutant, but not the WH/AA mutant, affected nucleosome and DNA binding (Figure S2A, D-H), likely because positively charged residues in Motif 3, mutated in 5A3, mediate non-specific DNA binding (the 5A1 and 5A2 mutants also bound nucleosomes; KW and AM, unpublished observations). In size-exclusion chromatography experiments with phosphorylated proteins, a direct interaction of ^GST^M18BP1^312-490^ with CENP-A nucleosomes was not visible, likely because of fast complex dissociation. Conversely, ^MBP^CENP-C^601-943^ bound robustly to CENP-A nucleosomes and to ^GST^M18BP1^312-490^ (Figure S4A-B). In line with the solid phase experiments, ^GST^M18BP1^312-490-5A1^ did not interact with ^MBP^CENP-C^601-943^, whereas ^GST^M18BP1^312-490-5A3^ and ^GST^M18BP1^312-490-WH/AA^ retained significant binding affinity (Figure S4C).

### The role of CENP-C

Thus, collectively, our binding analyses in vivo and in vitro indicate that Motif 1 and the SANTA domain are critical binding determinants to mitotic kinetochores. Motif 1 mediates an interaction with CENP-C^C^, while the main target of the SANTA domain does not seem to be present in our reconstitutions. Motif 3, while apparently dispensable in mitosis, contributes to kinetochore localization in interphase, likely by binding to CENP-HIKM or DNA. To corroborate the role of CENP-C in M18BP1 localization, we rapidly depleted CENP-C endogenously tagged with an auxin-inducible degron (Fachinetti et al., 2015) by addition of indole acetic acid (IAA) (Figure 4A-B). CENP-C depletion prevented kinetochore decoration of electroporated ^GST^M18BP1^312-490^ both in interphase and in mitosis, confirming that it is essential for recruitment of ^GST^M18BP1^312-490^. Importantly, we have previously shown that the localization of other CCAN subunits is not affected at early time points after rapid removal of CENP-C (Pesenti *et al*., 2022; Schweighofer et al., 2025), suggesting that CENP-C is directly involved in M18BP1 localization, and not indirectly through the stabilization of other CCAN subunits. The C-terminal nucleosome-binding motif of CENP-C (the CENP-C motif, Figure 1G) did not appear to be required for the interaction with M18BP1, as mutations in it did not affect the interaction with M18BP1 (Figure S5A). Rather, cross linking-mass spectrometry experiments were consistent with a putative interaction of ^GST^M18BP1^312-490^ with the Cupin domain (Figure S5C-D). Indeed, size-exclusion chromatography experiments demonstrated a robust interaction of the CENP-C Cupin domain (residue 775-943, lacking the nucleosome-binding motif) with ^GST^M18BP1^312-490^ (Figure S5B). AF predicted that Motif 1 of M18BP1 interacts with a pocket in the Cupin domain of CENP-C exposing the side chains of several conserved hydrophobic and positively charged residues (Figure 4C and Figure S5E-F). Combining mutations V858E, K880A, L890R, and F938A in this pocket (abbreviated as CENP-C^Cmut^, for <u>C</u>upin <u>mut</u>ant) entirely abrogated binding to ^GST^M18BP1^312-490^ (Figure 4D).

**Figure 4.**
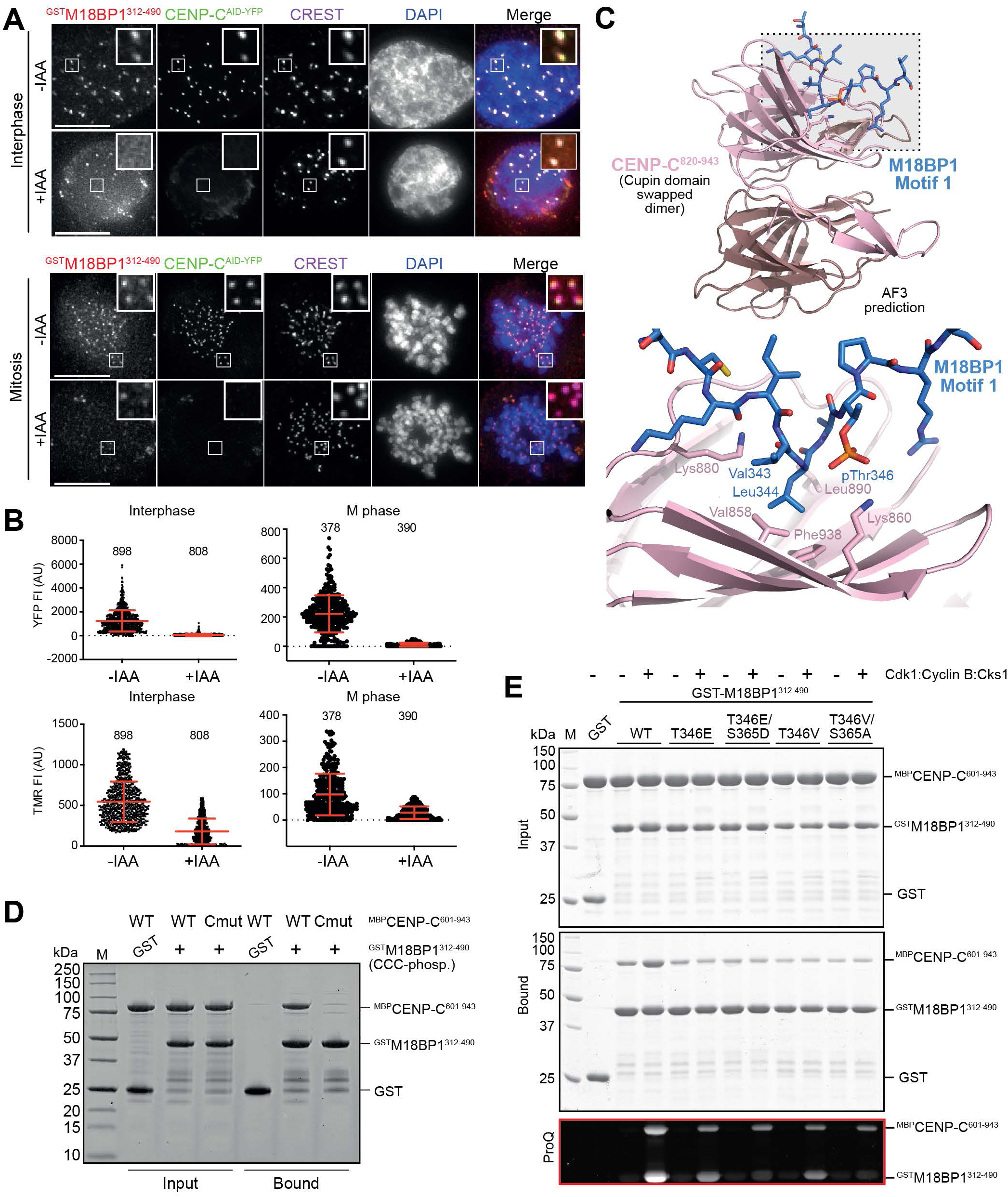
Rapid CENP-C depletion prevents efficient M18BP1 recruitment. (**A**) Representative images of fixed DLD-1 cells showing YFP fluorescence of endogenous CENP-C^AID-YFP^, TMR fluorescence of electroporated recombinant ^GST^M18BP1^312-490^ and immuno-stained α-tubulin in interphase (top) and mitosis (bottom). Centromeres were visualized by CREST sera, and DNA was stained by DAPI. Cells were treated with IAA as indicated to rapidly degrade endogenous CENP-C. White scale bars indicate 10 µm. (**B**) Quantification of the centromeric YFP-fluorescence intensities of endogenous CENP-C and the centromeric TMR-fluorescence intensities of the electroporated M18BP1 protein in interphase and mitosis. Centromeres were detected in the CREST channel using a script for semiautomated quantification. Red bars indicate mean and standard deviation of all quantified centromere fluorescence intensities. (**C**) AlphaFold prediction of the interaction of M18BP1 Motif 1 with the CENP-C Cupin domain. (**D**) GSH-resin pull-down assay with GST (negative control) and ^GST^M18BP1^312-490^ WT and Cmut (V858E, K880A, L890R, and F938A) variants as baits testing the binding to ^MBP^CENP-C^601-943^. The M18BP1 variants used as baits and, when applicable, the presence of CDK1:Cyclin B:Cks1 kinase complex are indicated above each lane. (**E**) GSH-resin pull-down assay with GST (negative control) and ^GST^M18BP1^312-490^ variants as baits testing the binding to ^MBP^CENP-C^601-943^. The M18BP1 variants used as baits and the addition of CDK1:Cyclin B:Cks1 kinase complex are indicated above each lane. ProQ staining demonstrating the phosphorylation intensities of the different ^GST^M18BP1^312-490^ variants and ^MBP^CENP-C^601-943^ in the bound fraction.

Thr346, at the boundary of motif 2, is situated in a Thr-Pro motif expected to be a substrate for proline-directed CDK activity. The AF prediction shows the phosphate to point towards Ly880 of the Cupin domain. Using ProQ staining, which visualizes phosphorylation, we detected reduced phosphorylation of the 5A2 mutant (Figure S1H), likely explaining why its binding to CENP-C appeared insensitive to phosphorylation (Figure 3F). Mutation of T346 to Val or Glu decreased phosphate incorporation by CDK1 to an even higher extent (Figure S1H). Mutation of M18BP1^Thr346^ to Glu did not create a phospho-mimetic mutant, as the levels of ^MBP^CENP-C^601-943^ bound to ^GST^M18BP1^312-490-T346E^ were no higher, and in fact slightly lower, than those observed with wild type ^GST^M18BP11^312-490^. Nonetheless, the mutation prevented enhanced binding of ^MBP^CENP-C^601-943^ in presence of CDK activity (Figure 4E). Essentially identical results were observed when Thr346 was mutated to Val, or after including an additional alanine mutation in another putative CDK site, Ser365, which removed phosphorylation of ^GST^M18BP1^312-490^ altogether (Figure 4E and Figure S1H).

### CENP-A deposition assays identify additional kinetochore-targeting region

^GFP^M18BP1^1-490^ contains the MIS18-binding region (Figure 5A) and localizes to kinetochores in G1 (Figure S1B, D). We therefore expected this construct to retain CENP-A loading capabilities in the absence of endogenous M18BP1. Confirming the efficiency of endogenous M18BP1 depletion, deposition of new CENP-A was entirely impaired in a cell line not expressing a ^GFP^M18BP1 rescue construct (Figure S6A-B). In agreement with our prediction, ^GFP^M18BP1^1-490^ promoted robust CENP-A deposition in early G1 phase in cells depleted of endogenous M18BP1 (Figure 5B-D; expression levels of rescue constructs are in Figure S1I). Conversely, all mutations affecting G1 kinetochore localization of ^GFP^M18BP1^1-490^, including the 5A1, 5A3, and WH/AA mutants, also prevented its ability to support CENP-A loading, confirming that centromere localization is essential for M18BP1’s function in CENP-A loading (Figure 5A-D). These mutants failed to localize also when expressed in the presence of endogenous M18BP1, and accordingly also displayed a dominant-negative effect on the ability of endogenous M18BP1 to load CENP-A (Figure S6C, quantified in Figure 5C-D). In contrast, the 5A2 mutant showed robust CENP-A loading in G1 (Figure S6D-F), in agreement with our observation that it lacks penetrance even when introduced in the minimal localization module ^GST^M18BP1^312-490-GFP^ (Figure 3B-D).

**Figure 5.**
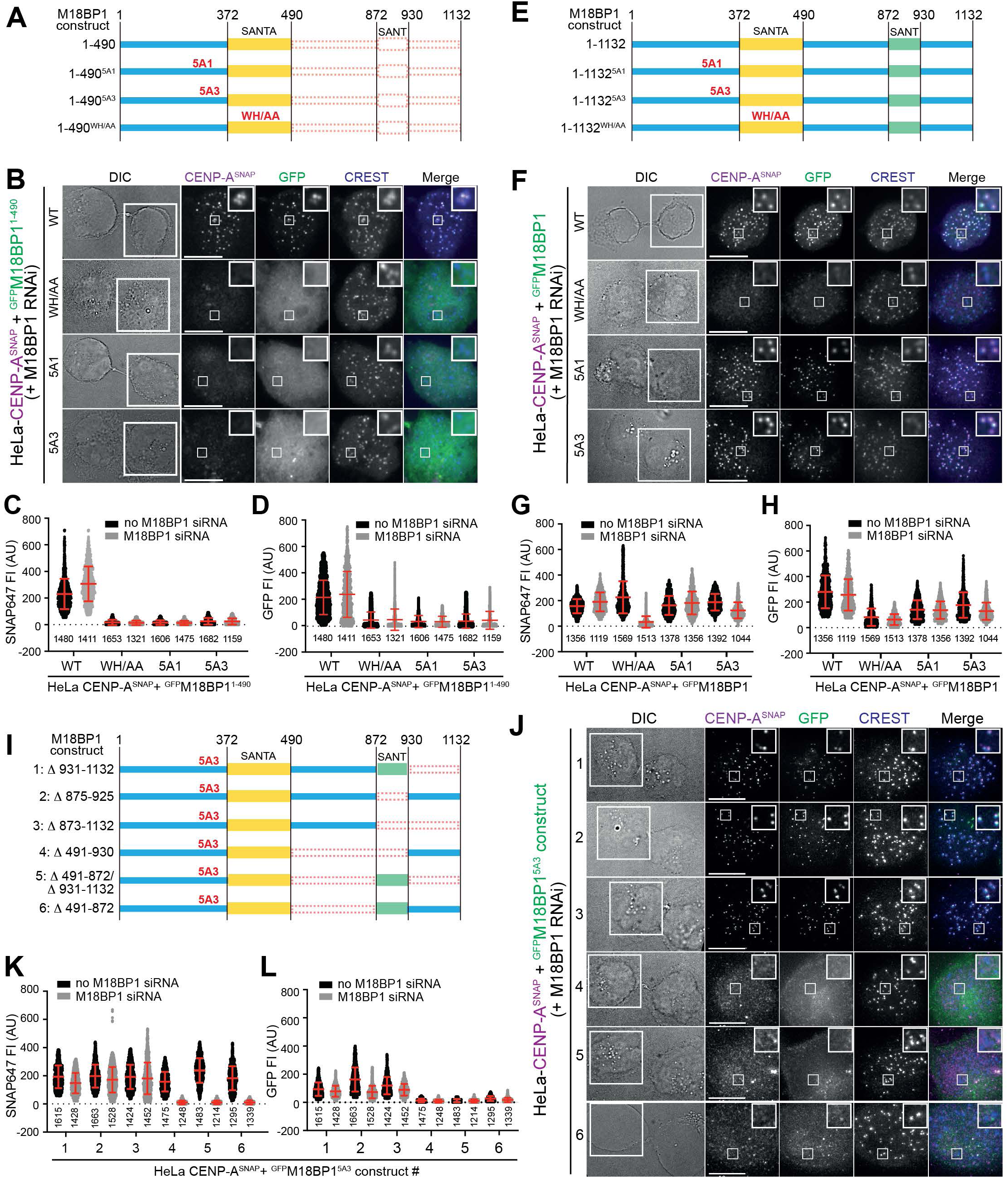
A C-Terminal region in M18BP1 provides further binding affinity to the centromere. (**A**) Schematic showing the expressed ^GFP^M18BP1^1-490^ variants in panel B (**B**) Representative images of fixed HeLa cells showing CENP-A^SNAP^ fluorescence labeled with SNAP-Cell 647-SiR and GFP fluorescence of indicated stably expressed M18BP1 variants in early G1. Centromeres were visualized by CREST sera. The G1 couple is shown in the differential interference contrast (DIC) channel. White scale bars indicate 10 µm. (**C**) and (**D**) Quantification of the centromeric CENP-A-SNAP and centromeric GFP fluorescence intensities, respectively, of HeLa cell lines stably expressing the indicated M18BP1 variants. Centromeres were detected in the CREST channel using a script for semiautomated quantification. Red bars indicate mean and standard deviation of all quantified centromere fluorescence intensities from three independent experiments. (**E**) Schematic showing the expressed ^GFP^M18BP1^1-1132^ variants in panel F (**F**) Representative images of fixed HeLa cells showing CENP-A^SNAP^ fluorescence labeled with SNAP-Cell 647-SiR and GFP fluorescence of indicated stably expressed M18BP1 variants in early G1. Centromeres were visualized by CREST sera. The G1 couple is shown in the differential interference contrast (DIC) channel. White scale bars indicate 10 µm. (**G**) and (**H**) Quantification of the centromeric CENP-A-SNAP and centromeric GFP fluorescence intensities, respectively, of HeLa cell lines stably expressing the indicated M18BP1 variants. Centromeres were detected in the CREST channel using a script for semiautomated quantification. Red bars indicate mean and standard deviation of all quantified centromere fluorescence intensities from three independent experiments. (**I**) Schematic showing the expressed ^GFP^M18BP1^5A3^ truncation variants in panel J (**J**) Representative images of fixed HeLa cells showing CENP-A^SNAP^ fluorescence labeled with SNAP-Cell 647-SiR and GFP fluorescence of indicated stably expressed M18BP1 variants in early G1. Centromeres were visualized by CREST sera. The G1 couple is shown in the differential interference contrast (DIC) channel. White scale bars indicate 10 µm. (**K**) and (**L**) Quantification of the centromeric CENP-A-SNAP and centromeric GFP fluorescence intensities, respectively, of HeLa cell lines stably expressing the indicated M18BP1 variants. Centromeres were detected in the CREST channel using a script for semiautomated quantification. Red bars indicate mean and standard deviation of all quantified centromere fluorescence intensities from three independent experiments. (**L**) Quantification of the centromeric GFP-fluorescence intensities of HeLa cell lines stably expressing the indicated M18BP1 variants. Centromeres were detected in the CREST channel using a script for semiautomated quantification. Red bars indicate mean and standard deviation of all quantified centromere fluorescence intensities from three independent experiments.

Next, we repeated these experiments with the same mutants introduced in full-length M18BP1 rather than M18BP1^1-490^ (Figure 5E; expression levels of rescue constructs are in Figure S1J). The detrimental effects of the 5A1 and 5A3 mutants on localization and CENP-A loading observed in the context of M18BP1^1-490^ were largely or completely rescued when the same mutations were expressed in the context of full-length M18BP1 (Figure 5F-H and Figure S7A). These observations suggest that regions downstream of residue 490 in the mutant full-length constructs contribute to kinetochore recruitment, thus rescuing kinetochore localization in presence of mutations that impair M18BP1^1-490^ localization. Only the WH/AA mutant in the SANTA domain, as well as a new deletion mutant where the entire SANTA domain had been deleted, were unable to localize to kinetochores and to load CENP-A in absence of endogenous M18BP1 (Figure 5F-H and Figure S7A-D). The SANTA deletion mutant, and to a minor extent the WH/AA mutant, had a dominant negative effect on CENP-A loading in presence of endogenous M18BP1, an effect that was more prominent in cells with high expression levels of these M18BP1 mutants (Figure S7E-G). These observations demonstrate that the SANTA domain is a crucial determinant of M18BP1 localization and CENP-A loading.

Next, we aimed to identify regions of M18BP1 downstream of residue 490 that contribute to kinetochore localization in the presence of the 5A3 mutation. For this, we fused various C-terminal fragments to a core segment encompassing residues 1-490 and carrying the 5A3 mutation (Figure 5I). We then asked which of these constructs localized correctly to kinetochores in G1 phase, thus rescuing the localization defect of M18BP1^1-490-5A3^ and if they loaded new CENP-A, in presence or absence of endogenous M18BP1. These experiments unequivocally identified residues 491-872 as being necessary and sufficient for rescuing the kinetochore recruitment and CENP-A loading deficiency when fused to M18BP1^1-490-5A3^ (while deletion of this segment did not result in a dominant-negative effect on CENP-A loading. Figure 5J-L and Figure S8A; Figure S1K shows expression levels of these constructs).

### Cell-cycle control of M18BP1 localization

The M18BP1 region comprised between 491 and 872 includes a previously identified CDK site, Thr653, that contributes to the suppression of mitotic localization of M18BP1 (Stankovic *et al*., 2017). As our analysis indicates that this residue is contained in a region contributing to kinetochore localization of M18BP1, we investigated its mechanism of action in mitosis and interphase. First, we generated ^GST^M18BP1^EGFP^ constructs encompassing the 312-490 or 312-872 fragments and carrying both the 5A1 and 5A3 mutations to completely inactivate the kinetochore localization determinants of the pre-SANTA region (Figure 6A and Figure S1L). Neither mutant localized to kinetochores in mitosis, but further mutating Thr653 to Val in the context of ^GST^M18BP1^312-872-5A1/5A3-EGFP^ rescued robust kinetochore localization (Figure 6B, D). As the region preceding the SANTA domain is impaired by the 5A1 and 5A3 mutations, this result is a strong indication that phosphorylation of Thr653 prevents kinetochore localization by suppressing a localization determinant within the 491-872 fragment. This determinant ought to be functional in interphase, when CDK activity is at its lowest levels. Accordingly, the ^GST^M18BP1^312-872-5A1/5A3-EGFP^ construct localized robustly to kinetochores in G1 phase, and the Thr653 mutant did not enhance its localization (Figure 6C, E).

**Figure 6.**
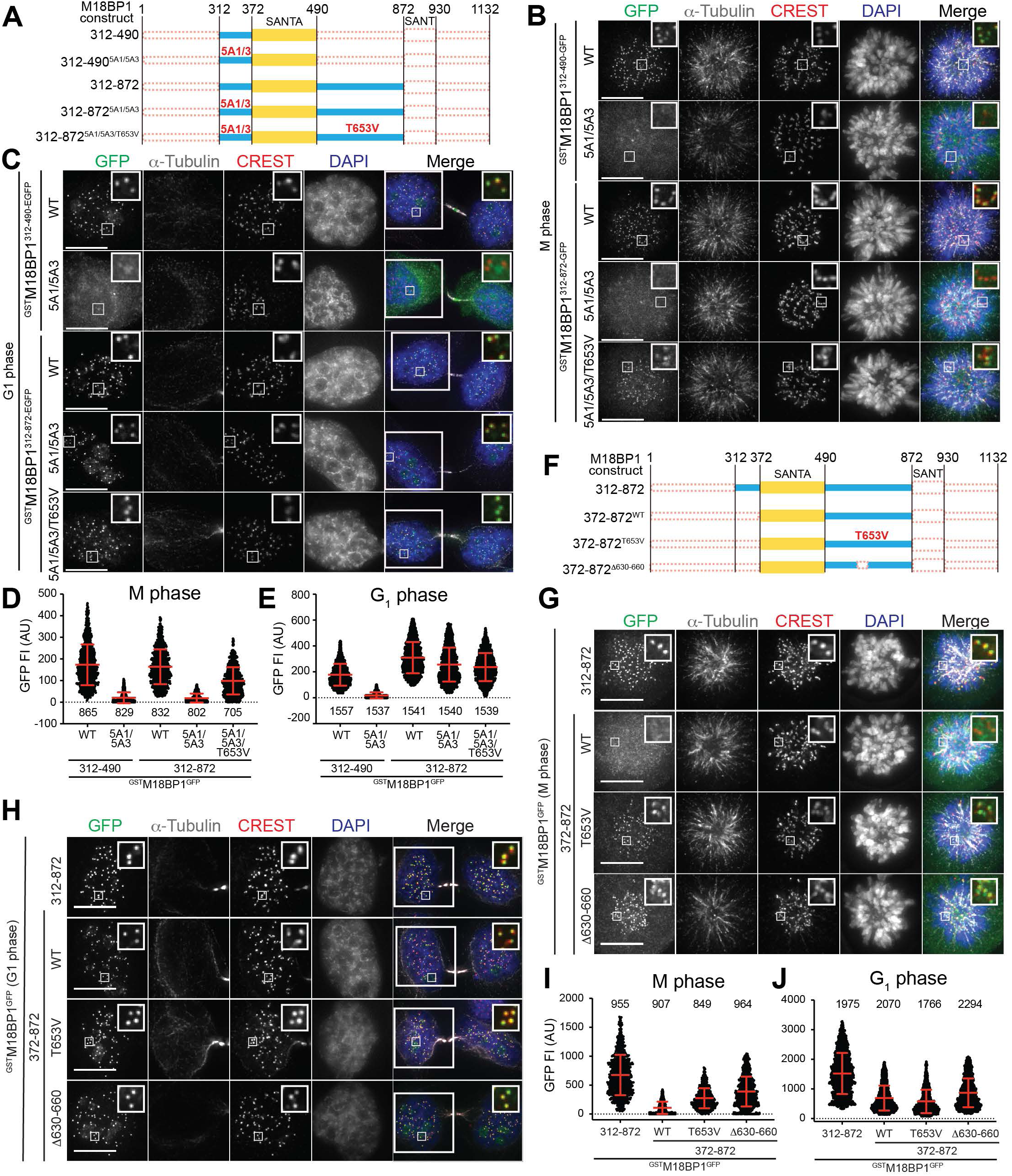
Thr653 is a regulatory key site of M18BP1 during mitosis. (**A**) Schematic showing the expressed ^GST^M18BP1^GFP^ variants in panels B and C (**B**) and (**C**) Representative images of fixed HeLa cells showing GFP fluorescence of indicated stably expressed M18BP1 variants and immuno-stained α-tubulin in mitosis and G1, respectively. Centromeres were visualized by CREST sera, and DNA was stained by DAPI. White scale bars indicate 10 µm. (**D**) and (**E**) Quantification of the centromeric GFP-fluorescence intensities of the indicated expressed M18BP1 variants in mitosis and G1, respectively. Centromeres were detected in the CREST channel using a script for semiautomated quantification. Red bars indicate mean and standard deviation of all quantified centromere fluorescence intensities from three independent experiments. (**F**) Schematic showing the expressed ^GST^M18BP1^GFP^ variants in panels G and H. (**G**) and (**H**) Representative images of fixed HeLa cells showing GFP fluorescence of indicated stably expressed M18BP1 variants and immuno-stained α-tubulin in mitosis and G1, respectively. Centromeres were visualized by CREST sera, and DNA was stained by DAPI. White scale bars indicate 10 µm. (**I**) and (**J**) Quantification of the centromeric GFP-fluorescence intensities of the indicated expressed M18BP1 variants in mitosis and G1, respectively. Centromeres were detected in the CREST channel using a script for semiautomated quantification. Red bars indicate mean and standard deviation of all quantified centromere fluorescence intensities from three independent experiments.

As a further demonstration that activation of the 491-872 segment requires dephosphorylation of Thr653 and that it can bypass a localization requirement for the pre-SANTA region of M18BP1, we generated new constructs lacking the pre-SANTA region altogether (Figure 6F). Unlike the control construct containing the pre-SANTA region, ^GST^M18BP1^372-872-EGFP^ failed to localize to mitotic kinetochores, but localized robustly to kinetochores in G1 phase, i.e. when Thr653 is expected to become dephosphorylated (Figure 6G-J). On the other hand, the ^GST^M18BP1^372-872-T653V-EGFP^ construct localized robustly to both mitotic and interphase kinetochores (Figure 6G-J). Thus, phosphorylation of T653 appears to control the function of a kinetochore-localization determinant in the 491-872 region. In the absence of the SANTA domain, this determinant was insufficient for kinetochore recruitment. The ^GST^M18BP1^491-872-GFP^ construct was unable to decorate kinetochores, regardless of the cell cycle phase and of whether the deletion was combined with the T653V mutation (Figure S8B-E).

We hypothesized that Thr653 may be part of a motif directly involved in the interaction with a centromere receptor, and that its phosphorylation may directly obstruct binding, a hypothesis reinforced by strong evolutionary sequence conservation around this site. We therefore deleted residues 630-660, comprising Thr653 and its surrounding region, and evaluated the localization of the resulting construct (^GST^M18BP1^372-872-Δ630-660-EGFP^) in mitosis and G1. Contrary to our hypothesis, ^GST^M18BP1^372-872-Δ630-660-EGFP^ localized to kinetochores in both interphase and mitosis, i.e. it had a localization pattern identical to that of the construct carrying the T653V mutation (Figure 6G-J and Figure S8F). These observations suggest that phosphorylation of Thr653 acts at a distance on M18BP1 localization, rather than in the immediate vicinity of the phosphorylation site.

### A crucial kinetochore-binding determinant in M18BP1^491-872^

N-terminally just adjacent to the 630-660 region of M18BP1 is a conserved sequence motif (residues 623-630) bearing sequence similarity to Motif 1, the CENP-C binding motif of M18BP1 examined above. A related sequence motif is also present in HJURP (Figure S5E). AF3 predicts this new motif, which we refer to as Motif 4, to bind the same pocket of the Cupin domain bound by Motif 1. *In vitro*, however, we only observed modest binding to the CENP-C Cupin domain of recombinant M18BP1 constructs encompassing Motif 4 (M18BP1^581-630^ or M18BP1^591-640^; LH and AM, unpublished observations). Nonetheless, we decided to test the effects of removing this sequence motif, which lies in a region predicted to be disordered (see AF prediction Q6P0N0). Like the wild type ^GST^M18BP1^372-872^ counterpart, ^GST^M18BP1^372-872^ carrying mutations in Motif 4 (residues 623-630) to alanine (^GST^M18BP1^372-872-623-630A-EGFP^) was not recruited to mitotic kinetochores. In interphase, there was residual recruitment of the mutant construct to chromatin foci, only a subset of which neighbored kinetochores but without overlapping with them (Figure 7A-D and Figure S8G). These observations indicate that Motif 4 encodes an additional determinant of kinetochore localization of M18BP1 in G1 phase.

**Figure 7.**
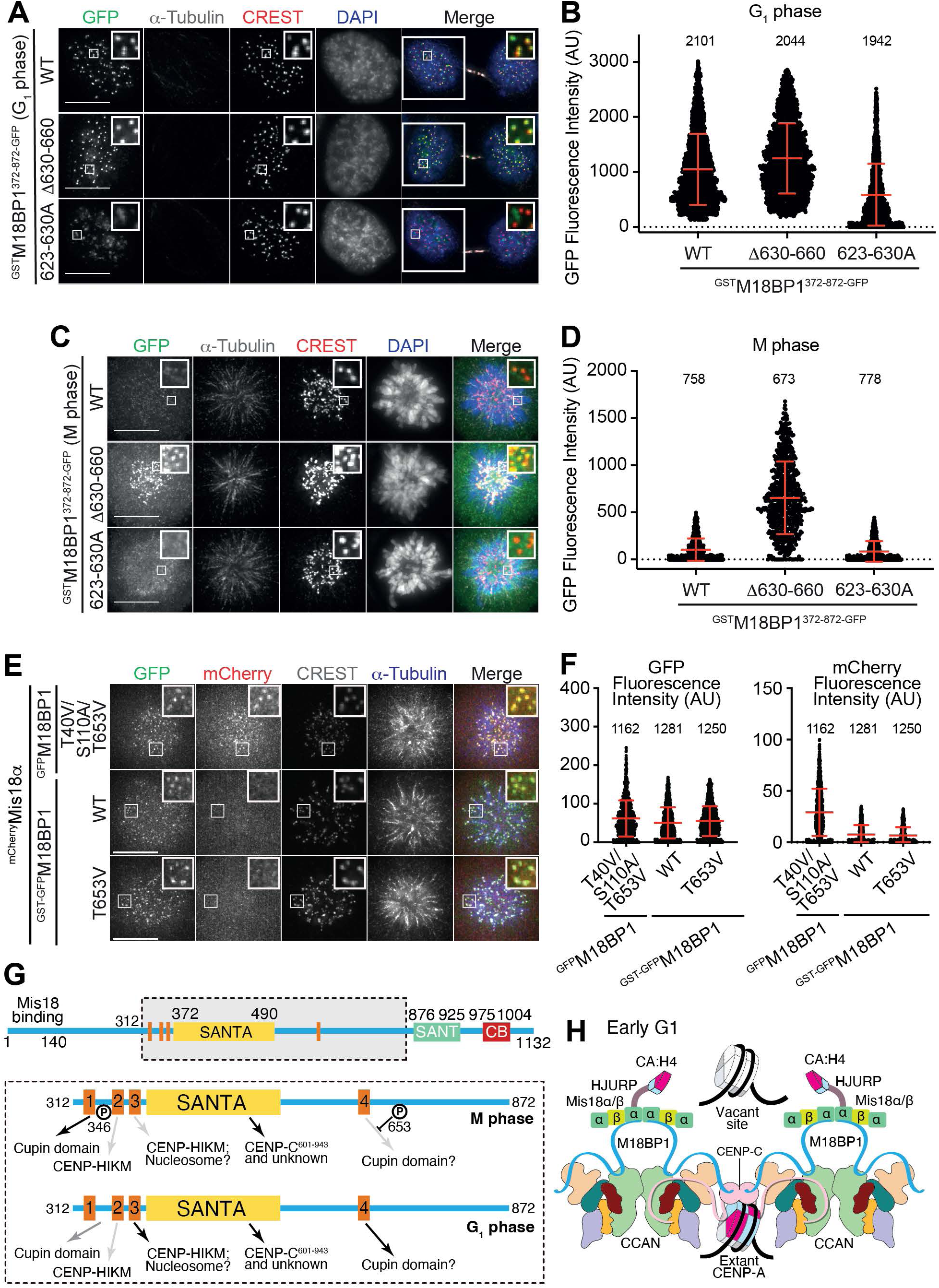
Crucial kinetochore-binding determinants in M18BP1 are sufficient for kinetochore localization upon dimerization. (**A**) Representative images of fixed HeLa cells showing GFP fluorescence of indicated stably expressed M18BP1 variants and immuno-stained α-tubulin in mitosis and G1. Centromeres were visualized by CREST sera, and DNA was stained by DAPI. White scale bars indicate 10 µm. (**B**) Quantification of the centromeric GFP-fluorescence intensities of the indicated expressed M18BP1 variants shown in panel **A**. Centromeres were detected in the CREST channel using a script for semiautomated quantification. Red bars indicate mean and standard deviation of all quantified centromere fluorescence intensities from three independent experiments. (**C**) Representative images of fixed HeLa cells showing GFP fluorescence of indicated stably expressed M18BP1 variants and immuno-stained α-tubulin in mitosis. Centromeres were visualized by CREST sera, and DNA was stained by DAPI. White scale bars indicate 10 µm. (**D**) Quantification of the centromeric GFP-fluorescence intensities of the indicated expressed M18BP1 variants shown in panel **C**. Centromeres were detected in the CREST channel using a script for semiautomated quantification. Red bars indicate mean and standard deviation of all quantified centromere fluorescence intensities from three independent experiments. (**E**) Representative images of fixed HeLa cells showing GFP fluorescence of indicated stably expressed M18BP1 variants, mCherry fluorescence of stably co-expressed Mis18α and immuno-stained α-tubulin in mitosis. Centromeres were visualized by CREST sera. White scale bars indicate 10 µm. (**F**) Quantification of the centromeric GFP and mCherry fluorescence intensities of the indicated expressed M18BP1 variants and co-expressed Mis18α shown in panel **E**. Centromeres were detected in the CREST channel using a script for semiautomated quantification. Red bars indicate mean and standard deviation of all quantified centromere fluorescence intensities from three independent experiments. (**G**) Model of M18BP1 recruitment to the centromere in M- and G1 phsae. The SANTA domain and four motifs mediate interactions to a variety of binding partners. The color of arrows reflects strength of binding (from black/strong to light grey/weak). (**H**) Role of dimerization in the recruitment of M18BP1 to CCAN and possible stoichiometries of CENP-A deposition by HJURP bound to the Mis18 complex.

### Determinants on M18BP1 are sufficient for kinetochore localization upon dimerization

To assess if the kinetochore-binding determinants of M18BP1 are sufficient for kinetochore recruitment after dimerization, we expressed ^GST-GFP^M18BP1 or ^GST-GFP^M18BP1^T653V^ in mitotic cells and ascertained that these constructs are recruited to kinetochores as expected (Figure 7E-F). As phosphorylation of T40 and S110 prevents the interaction of M18BP1 with the MIS18 complex in mitosis, ^mCherry^MIS18α co-expressed in the same cells was not identified at kinetochores (Figure 7E-F), demonstrating that the kinetochore localization determinants of M18BP1 are sufficient for kinetochore recruitment. Conversely, expression of ^GFP^M18BP1^T40V/S110A/T653V^, devoid of GST but allowing an ectopic interaction with the MIS18 complex in mitosis, led to substantial accumulation of co-expressed ^mCherry^MIS18α to mitotic kinetochores (Figure 7E-F), consistently with the idea that M18BP1 recruits the MIS18 complex when allowed to dimerize on it.

## Discussion

Unravelling the localization determinants of the CALM is key to decipher the molecular basis of epigenetic centromere inheritance. The work presented here on human M18BP1 rationalizes a large body of seemingly puzzling previous observations on the requirements for CALM localization. Previous work on M18BP1 had brought to light important species-specific features, for instance in nematodes (de Groot et al., 2021; Prosee et al., 2021). More widespread, a CENP-C-like motif present in M18BP1 orthologs in birds, reptiles, frogs, fish, and plants but not in mammals was shown to promote kinetochore localization of M18BP1 (French and Straight, 2019; French *et al*., 2017; Hori et al., 2017; Kral, 2015; Sandmann et al., 2017). The CENP-C-like motif of M18BP1, where present, appears to interact directly with CENP-A, like genuine CENP-C motifs (Jiang et al., 2023; Kato et al., 2013). Its deletion, however, only reduces M18BP1 kinetochore levels (French and Straight, 2019), suggesting the existence of additional localization determinants.

The identity of these additional determinants, and how mammalian M18BP1 replaces the CENP-C-like motif, has remained unclear. Deletions of the SANTA domain did not prevent M18BP1 localization in humans, chicken, or Arabidopsis (Hori *et al*., 2017; Lermontova et al., 2013; Stellfox *et al*., 2016). More recent work, however, identified a requirement for the SANTA domain for kinetochore localization of M18BP1 in humans and Xenopus (French and Straight, 2019; French *et al*., 2017). Our results provide new strong support to the essentiality of the SANTA domain in centromere recruitment, but also show the SANTA domain to be insufficient for kinetochore recruitment. We comment on this more extensively below. The SANT domain of mammalian M18BP1 has also been implicated in centromere localization (Dambacher *et al*., 2012). Here, we found no evidence supporting a major involvement of the SANT in M18BP1 localization, although we cannot exclude minor, non-essential roles.

Another aspect that had remained unclear is whether the localization determinants in M18BP1 are sufficient for kinetochore recruitment. M18BP1 localization to centromeres is contingent on the interaction with the MIS18α/β hexamer, which binds two M18BP1 molecules (Pan *et al*., 2017; Spiller *et al*., 2017; Subramanian et al., 2016; Thamkachy *et al*., 2024). By fusing M18BP1 to a strong constitutive dimer, we mimicked dimerization occurring when dephosphorylation of M18BP1 in G1 phase allows its interaction with MIS18α/β. A requirement for dimerization for robust kinetochore recruitment strongly suggests that the kinetochore target of M18BP1 is multivalent (as discussed below). Ectopic dimerization is sufficient for robust centromere recruitment of M18BP1 throughout the cell cycle, including mitosis, when CDK phosphorylation prevents M18BP1 from binding MIS18α/β. We observed localization of M18BP1 in absence of MIS18α/β, and also localization of M18BP1 constructs that do not bind MIS18α/β at all. Conversely, we did not observe localization of MIS18α/β under conditions that prevent its interaction with M18BP1. Thus, any contribution of the MIS18α/β to CALM localization (in addition to dimerizing M18BP1) may be dispensable, at least under the conditions of our assay (see Limitations of this study), as well as insufficient for MIS18α/β’s own centromere localization. In humans, the CENP-A-specific chaperone HJURP is recruited to MIS18α/β and interacts with CENP-C (Pan et al., 2019; Tachiwana *et al*., 2015; Thamkachy *et al*., 2024). Even HJURP recruitment, however, is exquisitely dependent on M18BP1, as it requires prior docking of PLK1 kinase onto two self-primed phosphorylation sites on M18BP1, Thr78 and Ser93 (Conti *et al*., 2024; Parashara *et al*., 2024). Thus, like MIS18α/β, also HJURP does not contain determinants sufficient for its kinetochore localization in absence of M18BP1. In essence, our work identified M18BP1, after dimerization, as the main reader of epigenetic centromere specification in humans.

Using the ectopic dimerization and localization assay in cells depleted of endogenous M18BP1, we identified the essential role of the M18BP1 SANTA domain. The apparent discrepancy with previous work referred to above likely stems from the fact that the previous experiments had been carried out in the presence of endogenous M18BP1, whose dimerization with exogenous mutant M18BP1 on the MIS18α/β might have led to substantial residual recruitment of mutant constructs. This interpretation is supported by our experiments involving overexpression of mutant M18BP1 lacking a functional SANTA domain. The SANTA domain was the only non-dispensable determinant of centromere localization of M18BP1 we have identified.

We determined the first experimental structure of the SANTA domain. This domain may be unique to M18BP1, as we could not identify close structural homologs in other proteins. A previous report identified CENP-C as a potential target of a construct consisting of pre-SANTA and the SANTA domain, and concluded CENP-C may be a direct target of the SANTA domain (French and Straight, 2019). We detected a very modest contribution of the SANTA domain to CENP-C binding, that was not affected by mutation of W413 and H414, two residues required for cellular localization of M18BP1. Given the relatively small size of the SANTA domain, and its convex binding interface, the target of the SANTA domain may be a motif embedded in an extended flexible segment of the polypeptide chain. As this target does not appear to be present in our biochemical reconstitutions of the CCAN or of centromeric nucleosomes, what could it be? Histone tails modified with centromere-specific modifications are potential candidates. However, enzymatic incorporation of two defining centromere-specific histone marks, Histone H4 Lys20 monomethylation or α-amino-trimethylation of the CENP-A N-terminus (H4-K20me1 or CENP-A-Nme3) (Bailey et al., 2016; Hori *et al*., 2014; Sathyan *et al*., 2017) did not increase the CENP-A-nucleosome-binding affinity for M18BP1 in vitro, nor was the binding sensitive to mutations of W413 and His414. KAT7, a histone acetyltransferase (HAT), binds M18BP1 and contributes to centromere stability (Ohzeki *et al*., 2016), and our future work will address the potential role of this enzyme in M18BP1 recruitment. While pre-nucleosomal CENP-A:H4 has also been shown to have specific post-translational modifications (Bailey *et al*., 2016; Sathyan *et al*., 2017; Shang et al., 2016), we doubt that they can function as guides for M18BP1, as the latter can be forced to localize in mitosis, where we expect no pre-nucleosomal CENP-A:H4 at centromeres (Jansen *et al*., 2007).

As the SANTA domain is required for kinetochore localization of M18BP1 both in interphase and in mitosis (upon forced dimerization), its elusive target exists at the centromere throughout the cell cycle. This target is a defining epigenetic marker of the centromere that remains elusive and whose identification is a clear goal of future studies. Even if essential, however, the SANTA domain is not sufficient for centromere localization and is complemented by adjacent pre- and post-SANTA disordered regions. Within these regions, we identify at least three distinct, biologically active centromere-targeting motifs, and a fourth motif also active in vitro (Figure 7G). Among the three biologically active motifs, two (Motifs 1 and 4) are putative CENP-C-binding motifs, while the other (Motif 3) binds at the interface of the CCAN subunits CENP-H and CENP-K but also interacts with nucleosomes in vitro, albeit non-specifically.

Phosphorylation of residues near these motifs influences their ability to interact with the centromere during different phases of the cell cycle. In the pre-SANTA region, Motif 1-3 mediate interactions with CCAN, including phosphorylation-dependent interactions of Motif 1 with the CENP-C Cupin domain, facilitated by phosphorylation of the pre-SANTA region at residue Thr346, and of Motif 3 with CENP-HIKM. During interphase, both the pre-SANTA and the post-SANTA become dephosphorylated, allowing the pre-SANTA and post-SANTA to switch roles as main drivers of kinetochore localization, as indicated by our observation that Motif 1 is dispensable for localization in interphase. The post-SANTA region in mammals may be functionally equivalent to the CENP-C-like binding motif in other organisms. Phosphorylation of Thr653 functionally limits this region during mitosis. In contrast, M18BP1 orthologs containing the CENP-C-like motif localize throughout the cell cycle in Xenopus, Caenorhabditis elegans, and chicken (Hori *et al*., 2017; Maddox *et al*., 2007; Moree *et al*., 2011; Perpelescu et al., 2015). While previous observations suggested that CENP-C may be dispensable for interphase localization of M18BP1 (French and Straight, 2019; French *et al*., 2017), our work in human cells clearly indicates that CENP-C is required also in interphase.

The cell-cycle dependence of kinetochore localization of M18BP1 is of special importance, not only in view of the role of this protein in CENP-A deposition, but also in view of the recently discovered function of M18BP1 in chromosome condensation through the loading of Condensin II, which is activated by CDK phosphorylation and engages a C-terminal region of M18BP1 apparently not involved in kinetochore recruitment (Borsellini et al., 2024; Wenda et al., 2021). During mitosis, when M18BP1 is phosphorylated by CDK1 and monomeric, and therefore only weakly associated to centromere, the pre-SANTA and SANTA act as localization determinants of M18BP1, and we speculate that these weak interactions may facilitate deposition of Condensin II. In summary, our observations delineate a defined hierarchy in kinetochore recruitment of the CALM, where M18BP1, after dimerization on MIS18α/β, is the only fully autonomous centromere targeting subunit of the human CENP-A loading machinery. Likely, the “epigenetic image” of M18BP1 is also a dimer built by CCAN and CENP-A nucleosomes in a specific reciprocal organization (Figure 7H). While recent structural work illuminated the organization of single CCAN complexes and a neighboring nucleosome (Pesenti *et al*., 2022; Tian et al., 2022; Yatskevich et al., 2022), our new observations on the mechanism of M18BP1 recruitment strongly suggest the existence of a higher order organization that acts as a multivalent target of the CALM. For instance, this may consist of neighboring CCAN complexes, possibly in different states of assemblage, flanking a CENP-A nucleosome. Ultimately, the crucial function of the loading machinery is to identify “temporary space” for stocking CENP-A between its deposition in G1 phase and its redistribution to the sister chromatids in S-phase. In this context, the local spatial organization of CCAN and of CENP-A nucleosomes at the target site is expected to be a crucial determinant of the outcome of the deposition reaction. Importantly, the MIS18α/β-M18BP1 octamer binds to a single HJURP molecule, which in turn binds a single CENP-A:H4 dimer (Barnhart *et al*., 2011; Pan *et al*., 2017; Thamkachy *et al*., 2024) (Figure 7H). These stoichiometries may hold key significance for stable centromere inheritance and their dissection is a key goal for future work.

## Limitations of this study

1) In a subset of our experiments, we score “functionality” of mutant M18BP1 in terms of ability to deposit CENP-A, assessed by fluorescence colocalization of newly deposited CENP-A with CREST. Even if this is a measure of functionality, we have no evidence that CENP-A is deposited at the correct place by these mutants. So, what we identify as being sufficient for CENP-A deposition may not always imply that CENP-A is correctly deposited. It may be deposited, but incorrectly, possibly with long term detrimental effects on centromere stability that we have not monitored. 2) Multivalency is a common strategy in nature for increasing binding affinity to a multivalent target. Many of our results are based on an artificial dimerization strategy designed to mimic the dimerization of M18BP1 on the MIS18α/β complex. We cannot rule out that this strategy, by artificially increasing the binding affinity of M18BP1 for an intrinsically multivalent target like the centromere, with multiple copies of CENP-A and associated CCAN, introduced artifacts rather than mimicking a consequence of MIS18α/β binding. We mitigated this concern by re-introducing many mutations or deletions into full-length, monomeric M18BP1.

## STAR ★Methods

### Key resource table

### Resource availability

#### Lead contact

Further information and requests for resources and reagents should be directed to Andrea Musacchio

#### Materials availability

All in-house-generated reagents described in this manuscript are available from the Lead Contact.

### Experimental model and subject details

#### Human cell lines

Parental Flp-In T-REx HeLa cells were a gift from S. Taylor (University of Manchester, Manchester, England, UK). Flp-In T-REx DLD-1–CENP-C–AID-EYFP cells were a gift from D. Fachinetti (Institut Curie, Paris, France) and D. C. Cleveland (University of California, San Diego, USA). These cells have both alleles of CENP-C tagged at the C terminus with an AID-EYFP fusion (Fachinetti *et al*., 2015). Furthermore, a gene encoding the plant E3 ubiquitin ligase osTIR1 was stably integrated into the genome of the cells. To induce rapid depletion of the endogenous AID-tagged CENP-C, 500 μM of the synthetic auxin Indole-3-acetic acid (IAA, Sigma Aldrich) was added to the cells. In all cell culture experiments, cells were cultured in Dulbecco’s modified Eagle’s medium (DMEM; PAN-Biotech) supplemented with 10% tetracycline-free fetal bovine serum (Thermo Fisher Scientific), 2 mM penicillin/streptomycin (PAN-Biotech), and 2 mM l-glutamine (PAN-Biotech) at 37°C in a 5% CO_2_ atmosphere. Expression of GFP-M18BP1 fusion proteins was induced by addition of doxycycline (50 ng/ml; Sigma-Aldrich) for at least 24 hours.

#### Bacterial strains

*E. coli* BL21CodonPlus(DE3)-RIL (Agilent Technologies, Santa Clara, California, United States) strains were cultured on LB agar or liquid media at 37 °C supplemented with ampicillin (100 µg/mL) to maintain the pETDuet or pGEX plasmids and with chloramphenicol (34 µg/mL) to maintain the extra copies of tRNA Genes in the CodonPlus strain.

### Method details

#### Plasmids and cloning

pGEX6PT-M18BP1^312-490^ (Conti *et al*., 2024) was modified to pGEX6PT-M18BP1^312-490^-LPETGG-6His by inserting the PCR-amplified sequence of M18BP1^312-490^-LPETGG-6His into the pGEX6PT backbone using the *Bam*HI and *Xho*I sites. The T346V/E, S365A/D, W413A, H414A and point mutations were introduced by PCR-based site-directed mutagenesis. The 5-Alanine substitution mutants (residues 341-345 mutated to Ala, named 5A1; residues 347-351 mutated to Ala, named 5A2; and residues 355-359 mutated to Ala, named 5A3, were all generated by inverse PCR. To clone pGEX6PT-M18BP1^312-371^-LPETGG-6His and pGEX6PT-M18BP1^372-490^-LPETGG-6His, the respective PCR amplified CDS of M18BP1 was inserted into the pGEX6PT-LPETGG-6His backbone using *Bam*HI and *Xho*I sites. pET-Duet-MBPT-SpyTag-CENP-C^601-943^-8His (Walstein *et al*., 2021) was used as template for the introduction of the CENP-C point mutations T734V, S773A, V858E, K880A, L890R, and F938A which were introduced by site-directed mutagenesis. pcDNA5-EGFP-M18BP1^1-1132^-P2AT2A-mCherry-Mis18α (Pan *et al*., 2017) was used as template to introduce the W413A, H414A, T653V, 5A1, 5A2 and 5A3 mutations as described above. The truncation mutants M18BP1^1-872-5A3^ and M18BP1^1-930-5A3^ were PCR amplified using pcDNA5-EGFP-M18BP1^1-1132-5A3^-2AT2A-mCherry-Mis18α as template and inserted using the BamHI and XhoI sites. The deletion mutants M18BP15^Δ875-925-5A3^, M18BP15^Δ491-930-5A3^, M18BP15^Δ491-872-5A3^, and M18BP15^Δ491-872-5A3^ were generated by PCR-based site-directed mutagenesis using pcDNA5-EGFP-M18BP1^1-1132-5A3^-P2AT2A-mCherry-Mis18α as template. pcDNA5-GST-M18BP1^312-490^-GFP was generated by inserting the CDS of GST-PresC-TEV and GFP into pcDNA5 using Gibson Assembly. The CDS of M18BP1^312-490^ was inserted using BamHI and XhoI sites. To generate pcDNA5-GST-M18BP1^312-872^-GFP, the CDS of M18BP1^312-872^ was inserted using BamHI and XhoI sites. pcDNA5-GST-M18BP1^312-872^-GFP was used as template to introduce the deletion mutant M18BP1^312-872Δ630-660^ and the alanine substitution mutant M18BP1^312-872^ ^623-660A^ using site-directed mutagenesis. To generate pcDNA5-LAP-M18BP1^1-490^-IRES, PCR-amplified EGFP-TEV-S-tag CDS was inserted into linearized pcDNA5-M18BP1^1-490^-IRES plasmid using Gibson assembly.

#### Protein expression and purification

The *E. coli* expression plasmid pETDuet-EGFP-M18BP1^372-490^-GNano-6His used for crystallization of the SANTA domain was used to transform BL21-CodonPlus(DE3)-RIL strain (Agilent Technologies, #230240). The transformed cells were cultured in TB medium to an OD_600_ of 0.8 and the protein expression was induced by adding IPTG to the final concentration of 0.5 mM and further incubation at 20 °C for 16 hours. Cells expressing EGFP-SANTA-GFPnanobody-6His was disrupted by Microfluidizer^®^ (Microfluidics) in a lysis buffer containing 50 mM HEPES pH 7.5, 300 mM NaCl, 1 mM DTE and 10 mM PMSF. The cleared lysate after centrifugation was applied to a Ni-affinity column and the bound protein was eluted using buffer containing 50 mM HEPES pH 7.5, 300 mM NaCl, 400 mM imidazole and 1 mM DTE. The eluate was applied to a Hiload 26/600 Superdex 200 pg SEC column (GE Healthcare) equilibrated with buffer containing 50 mM HEPES pH 7.5, 300 mM NaCl and 1 mM DTE. The elution fractions containing monomeric EGFP-SANTA-GFPnanobody-6His were pooled, concentrated, and stored at −80 °C.

The expression plasmids pGEXPT-M18BP1^312-490^-LPETGG-His (carrying the WT sequence or the 5A1, 5A2, 5A3, WH/AA, T346V, T346E, T346V/S365A, T346D/S365D mutations) were used to transform BL21-CodonPlus(DE3)-RIL strain. The transformed cells were cultured in TB medium to an OD_600_ of 0.8 and the protein expression was induced by adding IPTG to the final concentration of 0.2 mM and further incubation at 20 °C for 16 hours. Cells expressing GST-M18BP1^312-490^-LPETGG-His variants were lysed by sonication in a lysis buffer containing 50 mM HEPES pH 7.5, 500 mM NaCl, 10 % glycerol, 1 mM TCEP and 1 mM PMSF. The cleared lysate after centrifugation was incubated with cOmplete His-Tag purification resin (Roche) for 16 h at 4°C and the bound protein was eluted using lysis buffer supplemented with 400 mM imidazole. The concentrated eluate was applied to a HiLoad 16/600 Superdex 200 pg SEC column (Cytiva) equilibrated with buffer containing 20 mM HEPES pH 7.5, 300 mM NaCl, 2.5 % glycerol and 1 mM TCEP. The elution fractions containing the GST-M18BP1^312-490^-LPETGG-His variants were pooled, concentrated, and stored at −80 °C.

MBPT-SpyTag-CENP-C-601-943-His and HisTMBP-CENP-C1-600-SpyCatcher were expressed and purified as previously described (Walstein *et al*., 2021). MBPT-SpyTag-CENP-C-775-943-His and MBPT-SpyTag-CENP-C-601-943^T734V/S773A^-His and MBPT-SpyTag-CENP-C601-943_V858E,K880A,L890R,F938A-His point mutants were expressed and purified as MBPT-SpyTag-CENP-C-601-943-His.

The CCAN subcomplexes CENP-HIKM, CENP-LN, CENP-OPQUR and CENP-TWSX were expressed and purified as previously described (Pesenti et al., 2018; Pesenti *et al*., 2022; Walstein et al., 2021; Weir et al., 2016). The CENP-HI^Y488A^K^S113R, G181R, E182K, E185K, D186R, E214K, E217K^M mutant complex used in this study was expressed and purified exactly as the WT CENP-HIKM complex. The BPA-incorporated ^GST^M18BP1^312-490^-LPETGG-His variant was expressed in *E*. *coli* BL21(DE3) strain (Agilent Technologies, #200131). The pGEX plasmid encoding the M18BP1 gene with TAG codon at position 412 was used to transform *E*. *coli* cells together with pEVOL-pBpF. The cells were cultured in TB media supplemented with ampicillin, chloramphenicol and 0.2 % arabinose at 37 °C. Protein expression was induced by adding IPTG to the final concentration of 0.2 mM and the unnatural amino-acid BPA at a final concentration of 1 mM when OD_600_ of the culture reached 0.6. The culture was further incubated at 20 °C for 16 h. Purification of the GST-M18BP1^312-490^ ^412Bpa^-LPETGG-His variant was performed in the same way as the other GST-M18BP1^312-490^-LPETGG-His variants described above.

#### Sortase-mediated labeling of proteins with TMR-conjugated peptides

GST-M18BP1^312-490^-LPETGG was labeled with GGGGK peptides with a C-terminally conjugated Tetramethylrhodamine (TMR, Genscript) fluorophore using purified Sortase 7M mutant. Labeling was performed for ∼16 h at 4 °C by incubation of 20 µM MBP-M18BP1^312–490^-LPETGG with 200 µM GGGGK-TMR peptides and 1 µM Sortase 7M in the reaction buffer containing 20 mM HEPES (pH 7.5), 300 mM NaCl, 2.5 % glycerol and 1 mM TCEP. 7M Sortase and the excess of GGGGK-TMR peptides were removed by size-exclusion chromatography using a Superdex Increase 200 10/300 column.

#### Analytical Size-exclusion chromatography

Analytical size-exclusion chromatography of samples containing GST-M18BP1^312-490^ variants, CENP-HIKM, MBP-CENP-C^601-943^ and CENP-A^NCP^ was performed on a Superdex 200 Increase 5/150 column (Cytiva) in SEC buffer containing 20 mM HEPES (pH 7.5), 300 mM NaCl, 2.5% glycerol, and 1 mM TCEP on an ÄKTAmicro system. The proteins and CENP-A^NCP^ were incubated alone and in combination in a total volume of 40 µL SEC buffer. All proteins were diluted to 10 µM, whereas CENP-A^NCP^ was diluted to 5 µM. The samples were eluted under isocratic conditions at 4°C in SEC buffer at a flow rate of 0.15 ml/min. Elution of protein/NCP complexes was monitored at 280 nm and 254 nm wavelengths. 100 µL fractions were collected and analyzed by SDS-PAGE and Coomassie Brilliant Blue staining.

#### In vitro protein phosphorylation

When indicated, samples were phosphorylated using in-house generated CDK1:Cyclin-B:CKS1 (CCC). CCC was purified as described previously (Huis In ’t Veld *et al*., 2022). Phosphorylation reactions were set up in SEC buffer [20 mM HEPES (pH 7.5), 300 mM NaCl, 2.5 % glycerol and 1 mM TCEP] containing 50 nM CCC, 10 μM CENP-C or M18BP1 substrate, 2 mM adenosine triphosphate, and 10 mM MgCl_2_. Reaction mixtures were incubated at 30°C for 30 min, alternatively at 10 °C for 16 h.

#### Glutathione-resin pull-down assays

Glutathione-resin pull-down assays were performed to check the binding of nucleosomes and kinetochore components to GST-tagged M18BP1 truncation variants. The proteins were diluted with binding buffer [20 mM HEPES (pH 7.5), 300 mM NaCl, 2.5% glycerol 1 mM TCEP and 0.01 % Tween-20] to a final concentration of 5 µM in a total volume of 50 μl and mixed with 20 μl of Glutathione agarose beads (Cytiva/GE Healthcare). After mixing the proteins and the beads, 20 μl were taken as input. The rest of the solution was incubated at 4°C for 1 hour on a thermomixer (Eppendorf) set to 1200 rpm. To separate the proteins bound to the Glutathione beads from the unbound proteins, the samples were centrifuged at 800 *g* for 3 min at 4°C. The supernatant was removed, and the beads were washed four times with 500 μl of washing buffer (same composition as binding buffer, but NaCl was lowered from 300 mM to 150 mM). After the last washing step, 20 μl of 2× SDS-PAGE sample loading buffer was added to the dry beads. The samples were boiled for 5 min at 95°C and analyzed by SDS-PAGE and CBB (Coumassie Brilliant Blue) staining.

#### EMSA assays

DNA or NCPs alone or DNA/M18BP1 or NCP/M18BP1 complexes pre-incubated in SEC buffer supplemented with 0.01 % Tween-20 for 10 min on ice at indicated concentrations were run on a 5 % acrylamide gel. EMSA loading buffer (40 % sucrose, 0.025 % bromophenol blue) was added 1:5 to the samples before transferring them to the wells. The native PAGE was run in the cold room at 100 V constant voltage for 60 – 90 min using 1 X TB as running buffer. After the run, the DNA was stained using SYBR Gold (Thermo Fisher Scientific) according to the manufacturer’s instructions and subsequently the gel was scanned on a Chemidoc MP system (Bio-Rad).

#### DSBU Cross-linking mass spectrometry

Approximately 100 μg of GST-M18BP1^312-490^/CENP-HIKM or GST-M18BP1^312-490^/MBP-CENP-C^601-943^ complexes in cross-linking buffer [50 mM HEPES (pH 7.5), 300 mM NaCl, and 1 mM TCEP] were incubated with 3 mM DSBU (Disuccinimidyl Dibutyric Urea) (diluted from 200 mM stock solution in dimethyl sulfoxide; Alinda Chemical Limited) at 25°C for 1 h. The reaction was quenched by addition of 100 mM Tris-HCl (pH 8) and incubated for 30 min. Diagnostic SDS-PAGE gels of samples before and after the cross-linking reaction were run to ensure the completion of reaction. Samples were precipitated overnight with four volumes of cold acetone, spun down, and following the removal of the supernatant, the pellet was briefly air-dried before resuspension in urea. Samples were processed according to our previously described protocol (Pan et al., 2018) and analyzed using MeroX (Iacobucci et al., 2018). Proximity maps were visualized in Circos plots and arranged using Adobe Illustrator.

#### UV-induced crosslinking using Bpa

The Bpa-incorporated GST-M18BP1^312-490^-^412Bpa^ was diluted in 200 µL SEC buffer at 5 µM and was incubated with an equimolar amount of MBP-CENP-C-601-943. LED UV light with a wavelength of 365 nm (Nichia, NCSU276A) was used to irradiate the samples for 15 min on ice to activate the cross-linking reaction. The mixture was incubated with 50 µL GSH resin at 4 °C for 30 min to enrich the GST-tagged M18BP1 protein and the crosslinked adducts. The beads were washed four times with high salt washing buffer (containing 20 mM HEPES at pH 7.5, 500 mM NaCl and 1 mM TCEP). 25 µL dry beads were then used for on-beads digest, the other 25 µL beads were boiled for 5 min after addition of 50 µL 2 x SDS buffer. 10 µL and 30 µL were run in separate lanes on a gradient gel SDS PAGE. The gel was stained with Coumassie Blue for 1 h, destained overnight and used for in-gel recovery of the crosslinked species.

#### Generation of stable HeLa cell lines

Stable Flp-In T-REx HeLa cell lines constitutively expressing various GFP-tagged M18BP1 constructs were generated by Flp/FRT recombination. Deletion mutants and point mutants of M18BP1 were generated by PCR site-directed mutagenesis, and the sequences of all constructs were verified by Sanger sequencing (Microsynth Seqlab). M18BP1 constructs were cloned into a pcDNA5/FRT plasmid and co-transfected with pOG44 (Invitrogen), a plasmid expressing the Flp recombinase, into cells using X-tremeGENE (Roche) according to the manufacturer’s instructions. Following 2 weeks of selection in hygromycin B (250 μg/ml; Thermo Fisher Scientific) and blasticidin (4 μg/ml; Thermo Fisher Scientific), single-cell colonies were collected and subsequently expanded. Expression of the transgenes was checked by immunofluorescence microscopy and Western blot analysis.

#### RNAi transfection and CENP-A deposition experiment

Gene expression of endogenous M18BP1 was inhibited using a single small interfering RNA (siRNA) (sequence: 5’-GAAGUCUGGUGUUAGGAAAdTdT-3’, obtained from Thermo Fisher Scientific), which targets the coding region of M18BP1 mRNA. The expression of codon-optimized M18BP1 rescue constructs was not affected by the siRNA treatment. 30 nanomolars of M18BP1 siRNA was transfected using Lipofectamine RNAiMAX (Thermo Fisher Scientific) according to the manufacturer’s instructions. To induce the expression of GFP-tagged M18BP1 rescue constructs, doxycycline (50 ng/ml) (Sigma-Aldrich) was added to the cells at the time of siRNA transfection. Thymidine was added at 1 mM final concentration to arrest cells at G1/S transition. When cells were released from thymidine block, existing CENP-A-SNAP proteins were blocked using 10 µM SNAP-Cell Block (NEB) according to the manufacturer’s protocol. 10 µM STLC was used to arrest cells in prometaphase. The cells arrested in prometaphase were separated from other cells by mitotic-shake-off, released from STLC by three consecutive washing steps with 1 mL DMEM and placed in wells of 24-well plates containing poly-lysine coated coverslips. Three hours later, the cells in early G1 phase attached on the coverslips and were treated with 3 µM SNAP-Cell 647-SiR (NEB) to label newly synthesized SNAP-CENP-A according to the manufacturer’s protocol.

#### Protein electroporation

To electroporate recombinant TMR-labelled GST-M18BP1^312-490^ variants into either DLD-1– CENP-C–AID-EYFP cells or into HeLa cells, the Neon Transfection System Kit (Thermo Fisher Scientific) was used. Cells (3 × 10^6^) were trypsinized, washed with PBS, and resuspended in electroporation buffer R (Thermo Fisher Scientific) to a final volume of 90 μl. Recombinant protein was diluted 1:2 in buffer R to 50 μM, and 30 μl of the mixture was added to the 90-μl cell suspension. After mixing the sample, 100 μl of the mixture was loaded into a Neon pipette tip (Thermo Fisher Scientific) and electroporated by applying one 20 ms pulse with an amplitude of 1400 V. The sample was subsequently added to 15 ml of prewarmed PBS, centrifuged at 500 g for 3 min, and trypsinized for 7 min to remove noninternalized extracellular protein. After one additional PBS washing step and centrifugation, the cell pellet was resuspended in DMEM and transferred to a 12-well plate containing poly-l-Lysine–coated coverslips. After the electroporation procedure, DLD-1–CENP-C^AID-EYFP^ cells were additionally treated with 500 μM IAA (Sigma-Aldrich) to induce rapid depletion of endogenous CENP-C.

#### Immunofluoresence microscopy

Cells grown on poly-L-Lysine (0,0001 %, Sigma Aldrich) coated coverslips were fixed for 10 min at room temperature using a 4 % paraformaldehyde (PFA) solution diluted in phosphate-buffer saline (PBS, prepared in-house) followed by three washing steps with PPS. The PFA–fixed cells were permeabilized with PBS-T [PBS buffer containing 0.1 % Triton X-100] for 10 min and incubated with PBS-T containing 4 % bovine serum albumin (BSA) for 40 min. Cells were incubated for 90 min at room temperature with CREST/anticentromere sera (Antibodies Inc.; dilution 1:200 in 2 % BSA in PBS-T), washed three times with PBS-T, and were subsequently treated for 30 min with anti-human Alexa Fluor 647–conjugated secondary antibody (Jackson ImmunoResearch; dilution 1:200 in 2 % BSA in PBS-T). In some experiments, a-Tubulin (Sigma #T9026; dilution 1:4000 in 2 % BSA in PBS-T) or M18BP1 (generated in-house (Conti *et al*., 2024); dilution 1:500 in 2 % BSA in PBS-T) were additionally immuno-stained. To visualize DNA, 4′,6-diamidino-2-phenylindole (DAPI) (0.5 μg/ml; Serva) was added during the last washing step for 3 min. After drying, the coverslips were mounted with Mowiol mounting media (EMD Millipore) on glass slides.

#### Imaging and quantification of centromere fluorescence intensities

Fixed cell samples were imaged at room temperature using a 60x oil immersion objective lens on a DeltaVision deconvolution microscope. The DeltaVision Elite System (GE Healthcare, UK) is equipped with an IX71 inverted microscope (Olympus, Japan), a PLAPON ×60/1.42 numerical aperture objective (Olympus) and a pco.edge sCMOS camera (PCO-TECH Inc., USA). Images were acquired as 16 z-sections of 0.2 µM. The z-sections were converted into maximal intensity projections or average intensity projections, converted into 16-bit TIFF files and exported for further analysis. Quantification of centromere signals was performed using the software Fiji with a script for semiautomated processing as previously described (Pan *et al*., 2019; Walstein *et al*., 2021). Briefly, centromere spots were chosen based on the parameters of shape, size, and intensity using the images of the reference channel obtained with CREST staining, and their positions were recorded. In the images of the data channels, the mean intensity value of adjacent pixels of a centromere spot was subtracted as background intensity from the mean intensity value of the centromere spot. Intensity values were exported to Excel (Microsoft), the top and lowest 10% of data points were removed and the resulting values were plotted using GraphPad Prism software 9.0 (GraphPad Software).

#### Immunoblotting

HeLa cells were harvested by trypsinization, and the cell pellet was washed once with PBS. Cells were incubated in lysis buffer [75 mM HEPES (pH 7.5), 150 mM NaCl, 10 mM MgCl_2_, 10% glycerol, 0.1% NP-40, Benzonase (90 U/ml; Sigma-Aldrich), and protease inhibitor mix HP Plus (Serva)] for 30 min on ice. The lysed cells were centrifuged at 16,000*g* for 30 min at 4 °C, and SDS sample buffer was added to the supernatant. After Tricin–SDS-PAGE, the proteins were blotted on a nitrocellulose membrane using the Trans-Blot Turbo Transfer System (Bio-Rad) and were subsequently detected by Western blot analysis. The following primary antibodies were used: anti– α-Tubulin (1:8000; Sigma-Aldrich, T9026), anti-GFP (1:1000; generated in-house), and anti-Vinculin (1:10,000; Sigma-Aldrich V9131). As secondary antibodies, we used anti-mouse (1:10,000; Amersham NXA931), anti-rabbit (Cytiva(GE) NA934V) or anti-rat (1:10,000; Amersham NXA935) conjugated to horseradish peroxidase. After incubation with ECL Western blotting reagent (GE Healthcare), images were acquired with the ChemiDoc MP System (Bio-Rad) using Image Lab 5.1 software.

#### Crystallization of SANTA domain

The human M18BP1 SANTA domain (residues 372 to 490) was crystallized as the fusion protein construct EGFP-**AS**-SANTA-**LEGT**-anti-GFP Nanobody-GGHHHHHH (linker residues are shown in bold). The fusion protein was concentrated to 32 mg/ml in buffer containing 50 mM HEPES pH 7.5, 300 mM NaCl and 1 mM DTE. Crystals were grown by sitting-drop vapor-diffusion by mixing 1 µl protein solution with 1 µl reservoir solution containing 0.1 M MES pH 6.0, 50% PEG400, and 5 %w/v PEG3000 (optimized from JCSG core III screen (Qiagen, Hilden, Germany), condition F1) and incubation at 20 °C for 5–10 days. Crystals were fished directly from the sitting-drops and flash-frozen without addition of cryoprotectant.

#### Structure determination

Data was collected at 100 K using a Pilatus 6MF detector at the P11 beamline at the PETRA synchrotron in Hamburg, Germany, and integrated and scaled using XDS and XSCALE (Kabsch, 2010). The structure was solved in space group C2 with 2 molecules per asymmetric unit via molecular replacement with PHASER (Collaborative Computational Project, 1994) using the crystal structure of GFP:GFPnanobody complex (PDB code 3OGO) as template. Refinement with PHENIX (Adams et al., 2010) resulted in a model (Emsley et al., 2010) with good Ramachandran geometry and R_work_/R_free_ values (Table S1). The C-terminal residues GGHis_6_ were missing in the electron density for both molecules, as well as residues MV at the N-terminus of the second molecule. In addition, the “linker” between EGFP and the SANTA domain – encompassing the last 9 residues of EGFP and the first 7 (monomer B) or 8 (monomer A) residues of the SANTA domain) was disordered in both molecules. In the last rounds of refinement, the NCS constraints were removed, but apart from a relative rotation of the SANTA domains in monomers A and B by approx. 15° (relative to the EGFP/nanobody domains), no major conformational differences were apparent (overall main chain r.m.s.d difference 1.68Å for 445 C_α_ atoms, and 0.78 Å for 111 residues of the SANTA domains). The SANTA domain of monomer B was slightly less well defined compared to monomer A as indicated by increased B factors. The coordinates have been submitted to the protein data bank (PDB) with identifier 9R6H.

#### AlphaFold predictions

AF predictions were performed using AlphaFold2 (Jumper *et al*., 2021) (version 2.3.1) or AlphaFold3 (Abramson *et al*., 2024) (version 3.0.0, checked out from GitHub (https://github.com/google-deepmind/alphafold3 on January 12, 2025). For AlphaFold2, 10 models were predicted for the target protein complex comprising the CENP-C cupin domain (residues 820–943, comprising the most rigid part of the cupin domain) and M18BP1 (residues 325-265). The predictions were generated using the multimer pipeline to account for protein-protein interactions. For AlphaFold3, a total of 25 models were predicted using five different random seeds, with each seed generating five models, to enhance conformational sampling. This approach ensured diverse structural outputs for the same target complex. The models were evaluated based on two criteria: the overall Template Modeling Score (TM-score) and the local Distance Difference Test (LDDT) scores for the interface residues of the M18BP1 or HJURP peptide interacting with the CENP-C cupin domain (residues 820-943). LDDT scores were computed for specific interface residues to evaluate the accuracy of local interactions critical to the protein-ligand interface. For each method, the model with the highest overall TM-score was selected as the primary model. In cases where multiple models had comparable TM-scores, the model with the highest LDDT values for the interface residues was prioritized to ensure optimal representation of the interaction interface.

### Quantification and statistical analysis

Quantification of fluorescence intensities was performed as indicated above. Statistical analysis was performed with a rank sum, nonparametric test comparing two unpaired groups (Mann-Whitney test) in GraphPad Prism. Symbols indicate: n.s.= p > 0.05, * = p ≤ 0.05, ** p ≤ 0.01, *** = p ≤0.001 **** = p ≤0.0001.

## Acknowledgements

We thank the beamline staff of the beamline P11 at the PETRA synchrotron, Hamburg, Germany, for support, and our colleague R. Gasper-Schönenbrücher for help with data collection. We are grateful to D. Fachinetti D.C. Cleveland, and D. Foltz for sharing reagents; to T.-C. Li and D. Summerer for help setting up cell sorting experiments; to I. Hoffmann, C. Koerner, B. Voss, S. Wohlgemuth, T. Crocilla, M. Nasimi, M. Richter and M. Terbeck for general technical assistance and for preparation of CCAN subunits and protein kinases; to Duccio Conti for help with cell line preparation; and to F. Müller and P. Janning for help with mass spectrometry experiments.

## Author contributions

K.W.: Conceptualization, investigation, project administration, visualization, and writing (original draft preparation).

D.P.: Conceptualization, investigation, and visualization.

L.H.: Investigation, project administration, and visualization.

I.R.V.: Conceptualization, AlphaFold prediction, crystal structure refinement, and visualization.

A.M.: Conceptualization, funding acquisition, project administration, supervision, visualization, and writing (original draft preparation).

## Data and materials availability

All data needed to evaluate the conclusions in the paper are present in the paper and/or the Supplementary Materials. Additional data related to this paper may be requested from the authors.

## Supplemental Information

Document S1 containing Table S1 and Figures S1-S8

---

**Figure S1.**
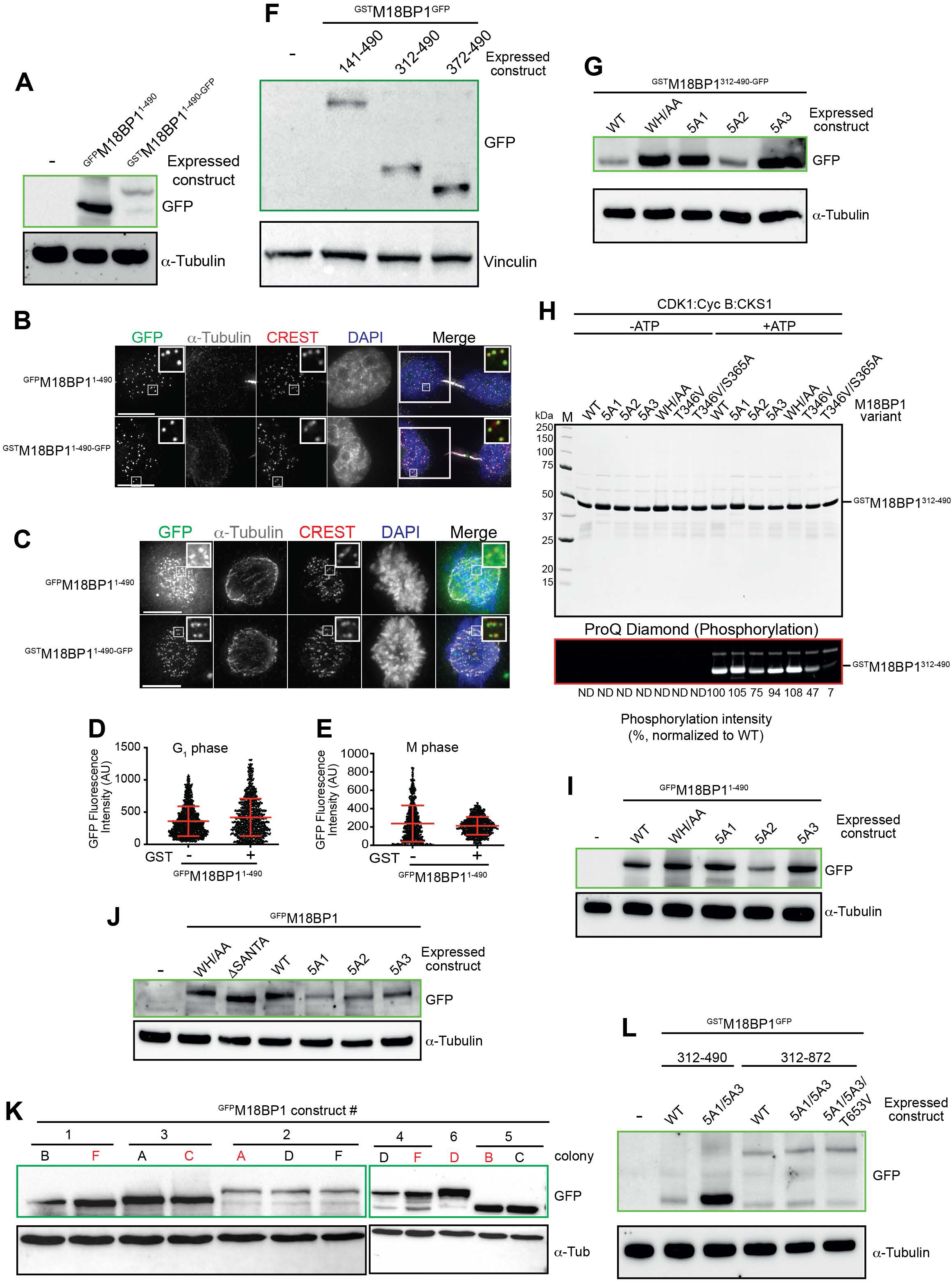
M18BP1 construct expression levels. (**A**) Anti-GFP and anti-α-Tubulin immunoblots of HeLa cell lysates expressing the indicated GFP tagged M18BP1 constructs. (**B**)-(**C**) Representative images of fixed HeLa cells showing GFP fluorescence of indicated stably expressed M18BP1 variants and immuno-stained α-tubulin in G1 and mitosis, respectively. Centromeres were visualized by CREST sera, and DNA was stained by DAPI. White scale bars indicate 10 µm. (**D**)-(**E**) Quantification of the centromeric GFP-fluorescence intensities of the indicated expressed M18BP1 variants in G1 and mitosis, respectively. Data shown as mean with SD. (**F**) Anti-GFP and anti-α-Vinculin immunoblots of HeLa cell lysates expressing the indicated ^GST^M18BP1^GFP^ constructs. (**G**) Anti-GFP and anti-Tubulin immunoblots of HeLa cell lysates expressing the indicated ^GST^M18BP1^312-490-GFP^ variants. (**H**) SDS-PAGE and ProQ staining showing the phosphorylation intensities of the indicated ^GST^M18BP1^312-490^ variants. All samples were incubated with CCC kinase, ATP was added as indicated. (**I**)-(**J**) Anti-GFP and anti-α-Tubulin immunoblots of HeLa cell lysates expressing the indicated GFP tagged M18BP1 constructs. (**K**) Anti-GFP and anti-α-Tubulin immunoblots of HeLa cell lysates from different colonies to compare the expression levels of the indicated GFP tagged M18BP1 constructs. The colonies marked in red were used for the experiment shown in figure 5 (I)-(L). (**L**) Anti-GFP and anti-α-Tubulin immunoblots of HeLa cell lysates expressing the indicated GFP tagged M18BP1 constructs.

**Figure S2.**
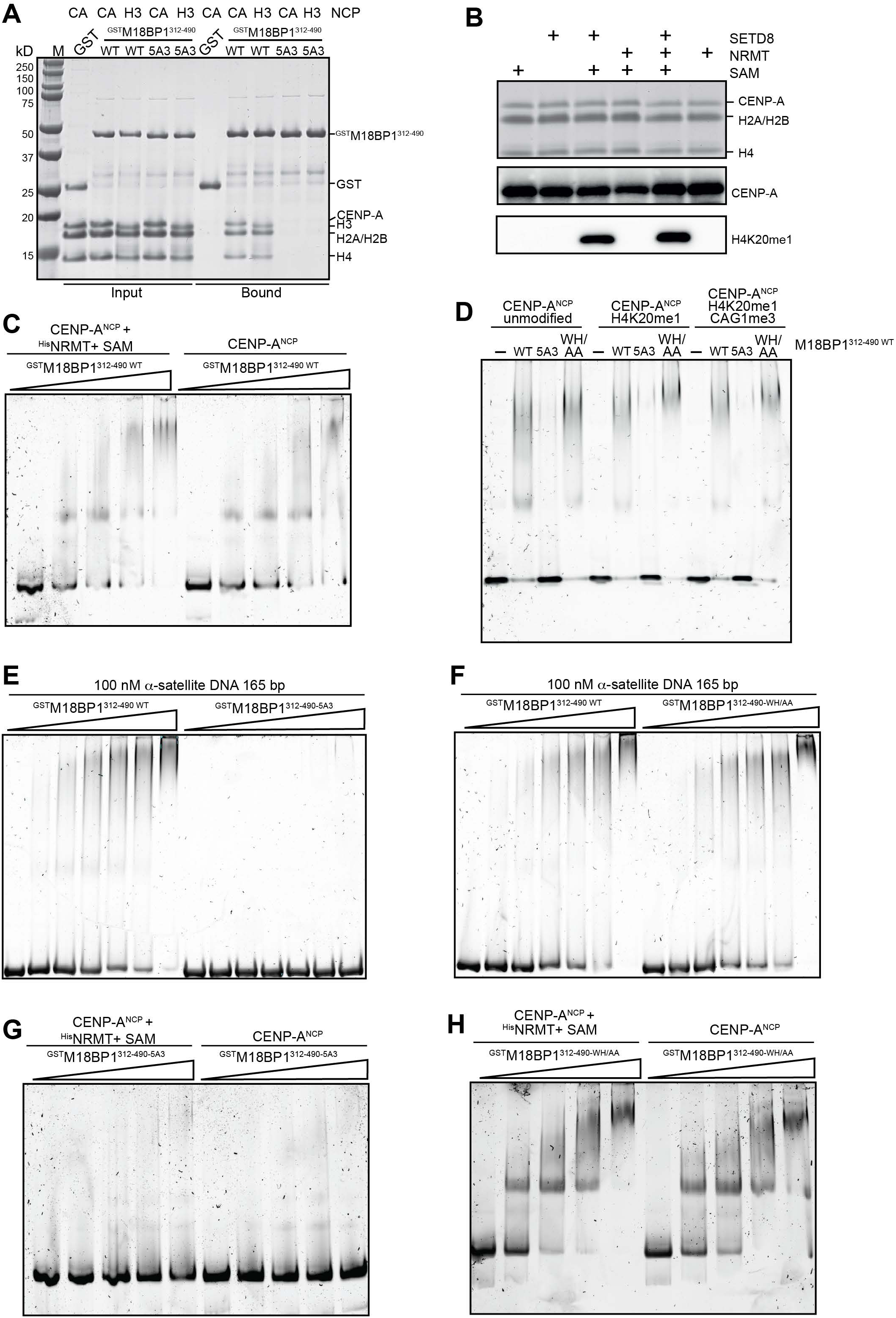
M18BP1 interacts with centromeric DNA and nucleosomes. (**A**) GSH-resin pull-down assay with GST (negative control) and ^GST^M18BP1^312-490^ as baits testing the binding to CENP-A^NCP^ (“CA”) and H3^NCP^. The NCP variant used as prey is indicated above each lane. (**B**) SDS-PAGE, anti-CENP-A and anti-H4K20me1 immunoblots of CENP-A^NCP^ treated with SETD8, NRMT and SAM as indicated above each lane. (**C**) EMSA experiment showing the interaction between ^GST^M18BP1^312-490^ and CENP-A^NCP^. The NCPs were added at 100 nM concentration, the M18BP1 concentration was varied from 0 mM (lowest concentration) to 1 mM (highest concentration). (**D**) EMSA experiment showing the interaction between ^GST^M18BP1^312-490^ variants and differently modified CENP-A^NCP^. The NCPs were added at 100 nM concentration, the indicated M18BP1 variants were added at 1 mM concentration. (**E**)-(**F**) EMSA experiments showing the interactions between a-satellite DNA and different ^GST^M18BP1^312-490^ variants. The DNA was added at 100 nM concentration, the indicated M18BP variant concentrations were varied from 0 mM (lowest concentration) to 2 mM (highest concentration). (**G**)-(**H**) EMSA experiments showing the interactions between ^GST^M18BP1^312-490^ variants and CENP-A^NCP^. The NCPs were added at 100 nM concentration, the M18BP1 concentration was varied from 0 mM (lowest concentration) to 1 mM (highest concentration).

**Figure S3.**
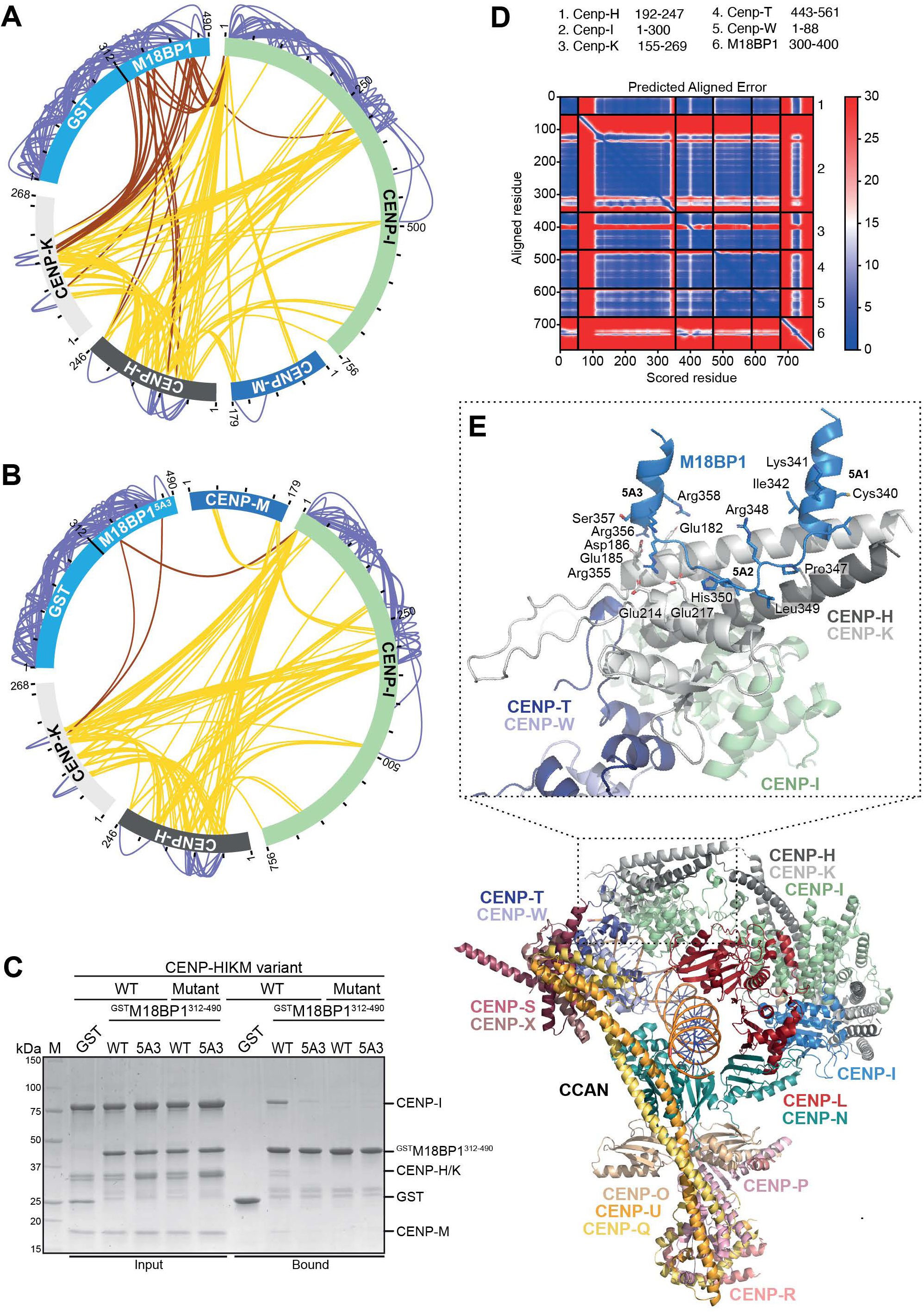
The interaction between M18BP1 and CENP-HIKM. (**A**) and (**B**) Schematic showing the results of DSBU crosslinking experiments followed by mass spectrometry analysis. In (A) ^GST^M18BP1^312-490^ ^WT^ was incubated with CENP-HIKM complex. In (B) ^GST^M18BP1^312-490^ ^5A3^ was incubated with CENP-HIKM complex. Intermolecular crosslinks are highlighted in red, intramolecular crosslinks are highlighted in yellow. (C) GSH-resin pull-down assay with GST (negative control) and ^GST^M18BP1^312-490^ (WT and 5A3 mutant) as baits testing the binding to WT and Mutant CENP-HIKM complexes. The M18BP1 variants used as baits and the CENP-HIKM variants used as preys are indicated above each lane. (**D**) and (**E**) AlphaFold3 prediction of the interaction of M18BP1 residues 300-400 with the CENP-HIKM complex. (D) shows the PAE and in (E) a cartoon model of the interaction in the context of the full CCAN is depicted.

**Figure S4.**
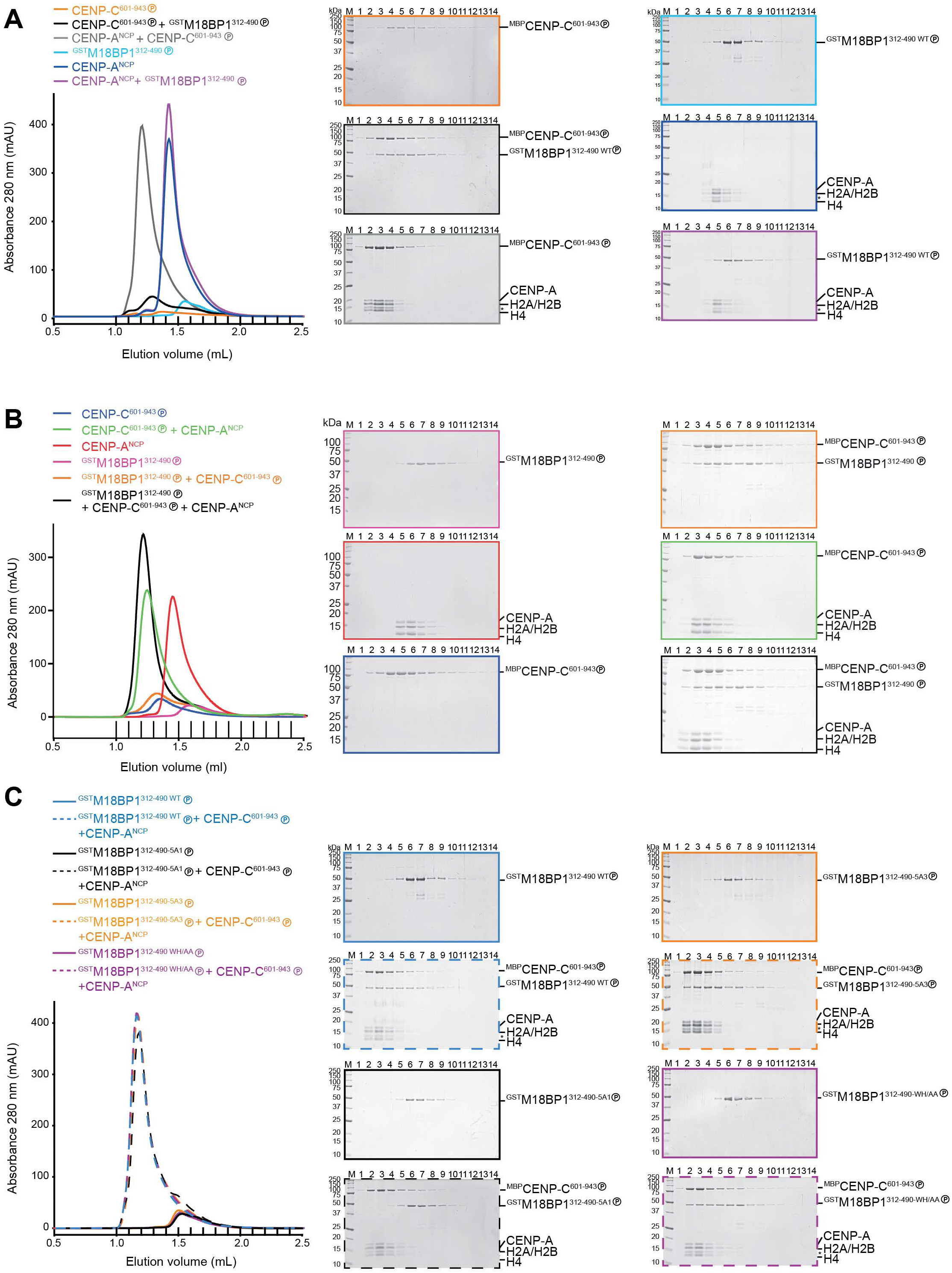
The interaction between M18BP1 and CENP-A is mediated by CENP-C. (**A**)-(**C**) Chromatograms and SDS PAGEs of analytical size exclusion (SEC) experiments. ^MBP^CENP-C^601-943^ and ^GST^M18BP1^312-490^ variants were incubated as indicated at 10 µM, CENP-A^NCP^ at 5 µM for 30 min. The individual proteins or incubated mixtures were loaded on an analytical Superdex 200 column. The 14 indicated fractions between 1 mL and 2.5 mL elution volume were subsequently analyzed by SDS PAGE. Components pre-phosphorylated by CCC kinase are marked by an encircled P.

**Figure S5.**
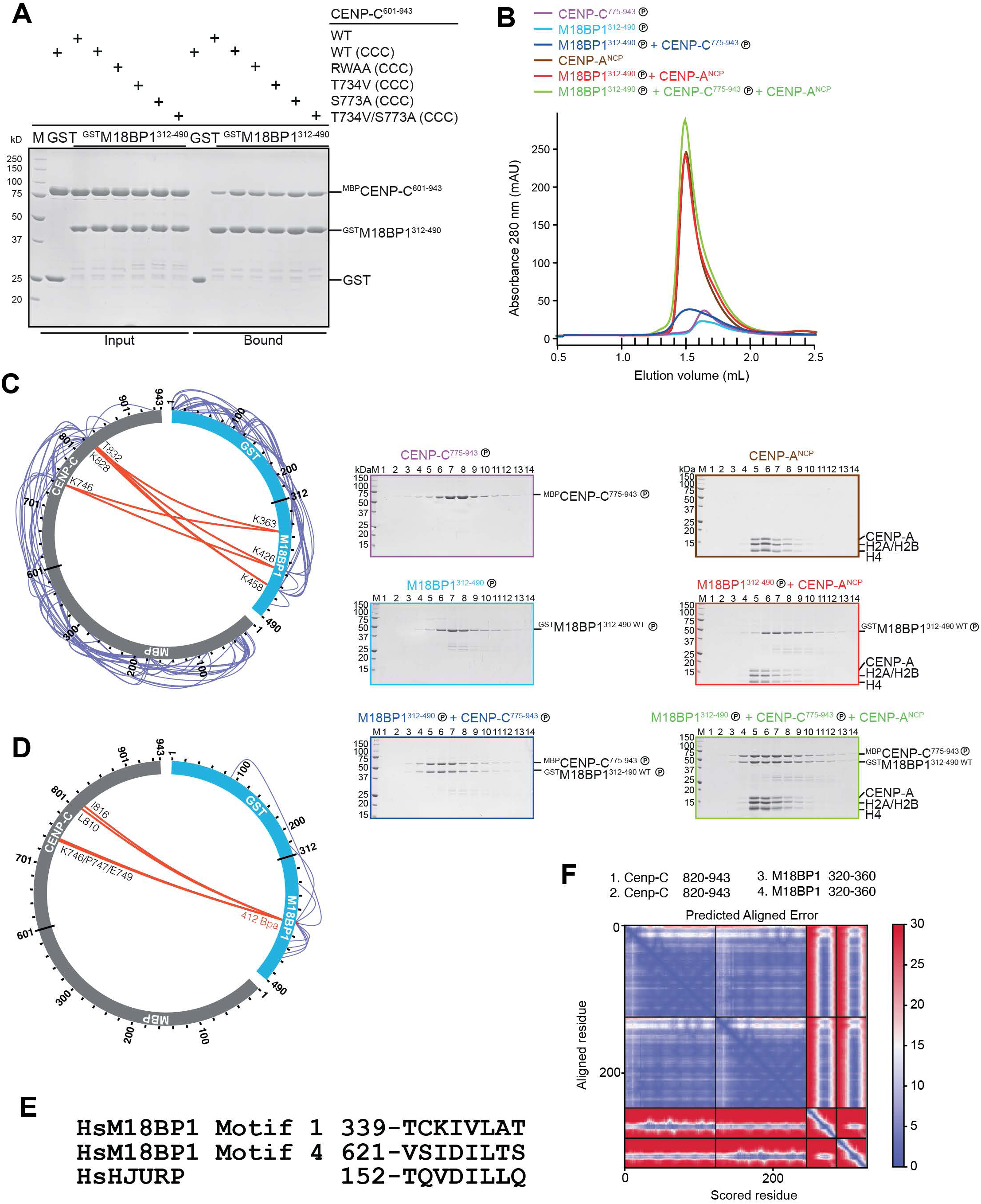
The interaction between M18BP1 and *CENP-C*. (**A**) GSH-resin pull-down assay with GST (negative control) and ^GST^M18BP1^312-490^ variants as baits testing the binding to different ^MBP^CENP-C^601-943^ constructs carrying mutations in conserved residues of the CENP-C motif, known to bind CENP-A nucleosomes. The M18BP1 variants used as baits and the addition of CDK1/Cyclin B/CKS1 (CCC) kinase complex are indicated above each lane. (**B**) Chromatograms and SDS PAGEs of analytical size exclusion (SEC) experiments. ^MBP^CENP-C^775-943^ and ^GST^M18BP1^312-490^ were incubated as indicated at 10 µM, CENP-A^NCP^ at 5 µM for 30 min. The individual proteins or incubated mixtures were loaded on an analytical Superdex 200 column. The 14 indicated fractions between 1 mL and 2.5 mL elution volume were subsequently analyzed by SDS PAGE. Components pre-phosphorylated by CCC kinase are marked by an encircled P. (**C**) Schematic showing the results of a DSBU crosslinking experiment followed by mass spectrometry analysis. ^GST^M18BP1^312-490WT^ was incubated with ^MBP^CENP-C^601-943^. Intermolecular crosslinks are highlighted in red, intramolecular crosslinks are highlighted in blue. Crosslinks involving the GST or MBP tags were excluded for clarity. (**D**) Schematic showing the results of UV-induced crosslinking experiments followed by mass spectrometry analysis. Ultraviolet (UV) light was used to induce crosslinking of CENP-C^601-943^ to ^GST^M18BP1^312-490_412BPA^(Benzoyl-phenylalanine introduced at residue 412). Intermolecular crosslinks are highlighted in red, intramolecular crosslinks are highlighted in blue. (E) Similarity of motifs in M18BP1 (Motifs 1 and 4) and HJURP. (**F**) AlphaFold3 prediction of the interaction of M18BP1 residues 300-400 with the CENP-HIKM complex. The PAE plot is shown.

**Figure S6.**
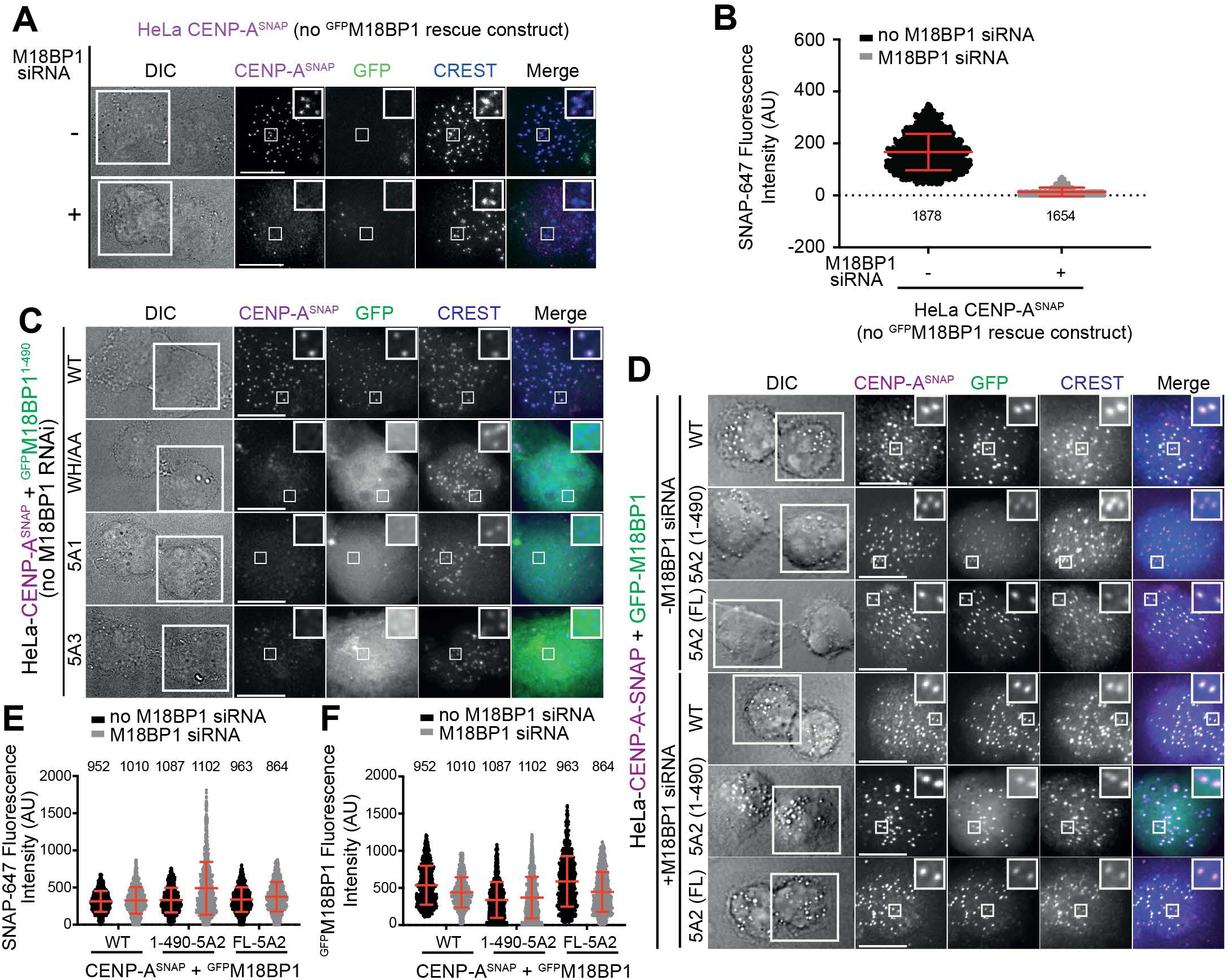
M18BP1^1-490^ does not tolerate 5A1, 5A3 and WH/AA mutations. (**A**) Representative images of fixed HeLa cells showing CENP-A^SNAP^ fluorescence labeled with SNAP-Cell 647-SiR and GFP autofluorescence in the absence of a ^GFP^M18BP1 rescue construct in early G1 in the absence or presence of M18BP1 siRNA treatment. Centromeres were visualized by CREST sera. The G1 couple is shown in the differential interference contrast (DIC) channel. White scale bars indicate 10 µm. (**B**) Quantification of the centromeric CENP-A-SNAP-fluorescence of a HeLa cell line that does not express any ^GFP^M18BP1 construct in the absence or presence of M18BP1 siRNA treatment. Centromeres were detected in the CREST channel using a script for semiautomated quantification. Data shown as mean with SD. (**C**) Representative images of fixed HeLa cells showing CENP-A^SNAP^ fluorescence labeled with SNAP-Cell 647-SiR and GFP fluorescence of indicated stably expressed M18BP1 variants in early G1. Centromeres were visualized by CREST sera. The G1 couple is shown in the differential interference contrast (DIC) channel. White scale bars indicate 10 µm. (**D**) Representative images of fixed HeLa cells showing CENP-A^SNAP^ fluorescence labeled with SNAP-Cell 647-SiR and GFP fluorescence of indicated stably expressed M18BP1 variants in early G1 in the absence or presence of M18BP1 siRNA treatment. Centromeres were visualized by CREST sera. The G1 couple is shown in the differential interference contrast (DIC) channel. White scale bars indicate 10 µm. (**E**) – (**F**) Quantification of the centromeric CENP-A-SNAP-fluorescence and centromeric GFP-fluorescence intensities, respectively, of HeLa cell lines stably expressing the indicated M18BP1 variants in the absence or presence of M18BP1 siRNA treatment. Centromeres were detected in the CREST channel using a script for semiautomated quantification. Data shown as mean with SD.

**Figure S7.**
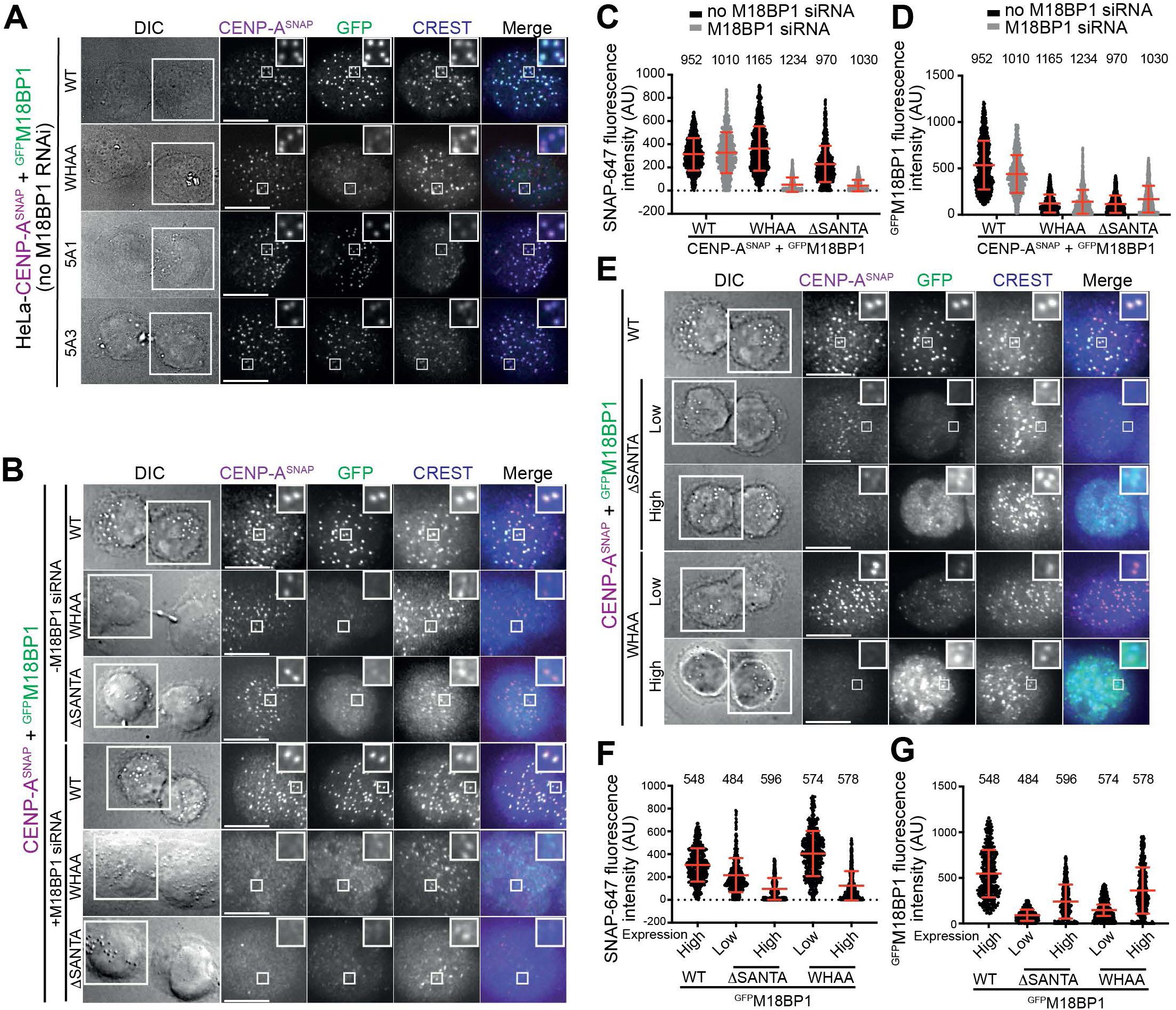
Additional CENP-A Loading experiments. (**A**) Representative images of fixed HeLa cells showing CENP-A^SNAP^ fluorescence labeled with SNAP-Cell 647-SiR and GFP fluorescence of indicated stably expressed M18BP1 variants in early G1. Centromeres were visualized by CREST sera. The G1 couple is shown in the differential interference contrast (DIC) channel. White scale bars indicate 10 µm. (**B**) Representative images of fixed HeLa cells showing CENP-A^SNAP^ fluorescence labeled with SNAP-Cell 647-SiR and GFP fluorescence of indicated stably expressed M18BP1 variants in early G1 in the absence and presence of M18BP1 siRNA treatment. Centromeres were visualized by CREST sera. The G1 couple is shown in the differential interference contrast (DIC) channel. White scale bars indicate 10 µm. (**C**)-(**D**) Quantification of the centromeric CENP-A-SNAP and GFP fluorescence intensities, respectively, of HeLa cell lines stably expressing the indicated M18BP1 variants in the absence and presence of M18BP1 siRNA treatment. Centromeres were detected in the CREST channel using a script for semiautomated quantification. Data shown as mean with SD. (**E**) Representative images of fixed HeLa cells showing CENP-A^SNAP^ fluorescence labeled with SNAP-Cell 647-SiR and GFP fluorescence of indicated stably expressed M18BP1 variants in early G1. Centromeres were visualized by CREST sera. The G1 couple is shown in the differential interference contrast (DIC) channel. White scale bars indicate 10 µm. (**F**)-(**G**) Quantification of the centromeric CENP-A-SNAP and GFP fluorescence intensities of HeLa cell lines stably expressing the indicated M18BP1 variants. Centromeres were detected in the CREST channel using a script for semiautomated quantification. Data shown as mean with SD.

**Figure S8.**
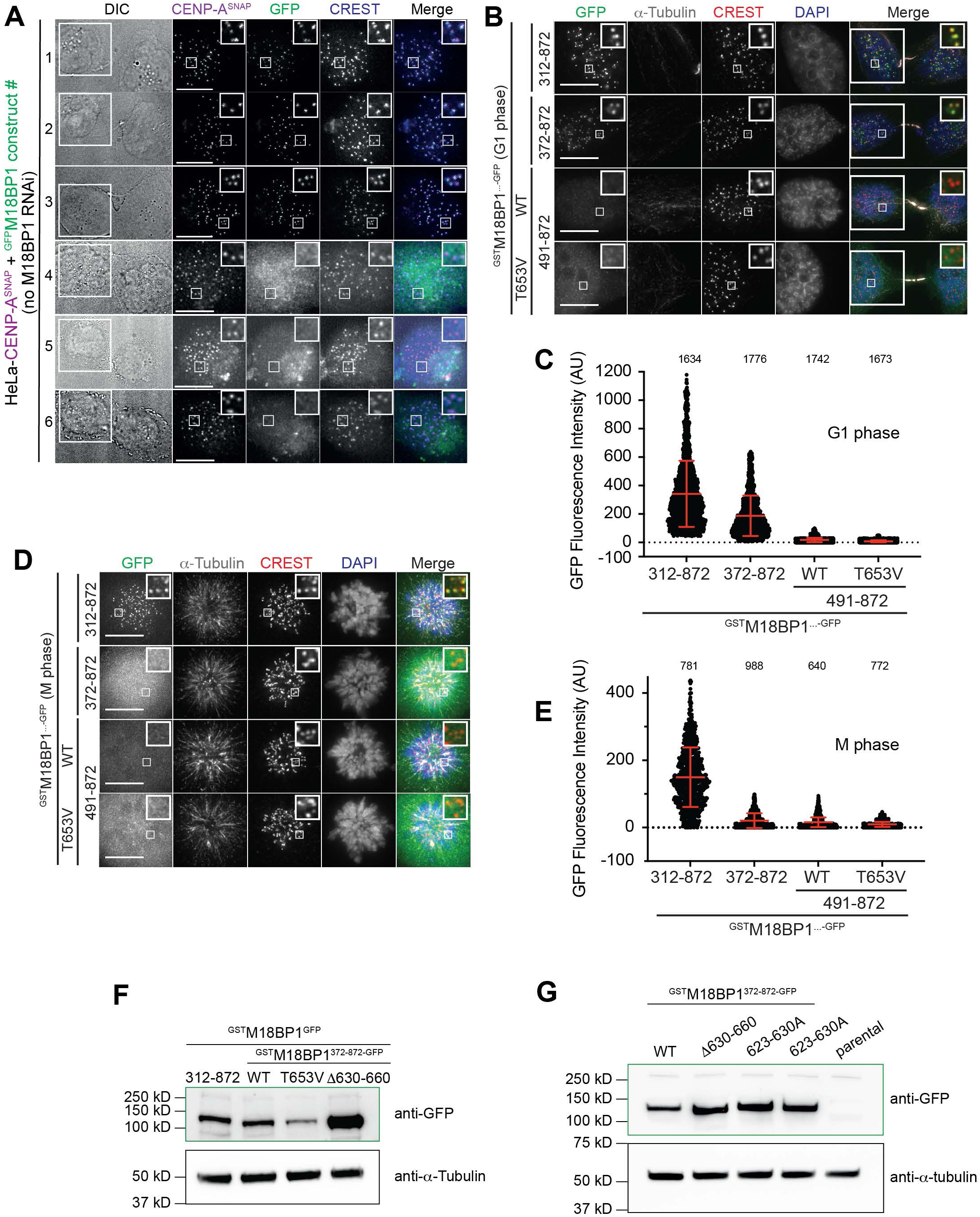
The kinetochore-localization determinant of the 491-872 region depends on the presence of the SANTA domain. (**A**) Representative images of fixed HeLa cells showing CENP-A^SNAP^ fluorescence labeled with SNAP-Cell 647-SiR and GFP fluorescence of indicated stably expressed M18BP1 variants in early G1. Centromeres were visualized by CREST sera. The G1 couple is shown in the differential interference contrast (DIC) channel. White scale bars indicate 10 µm. (**B**) Representative images of fixed HeLa cells showing GFP fluorescence of indicated stably expressed ^GST^M18BP1^GFP^ variants and immuno-stained α-tubulin in G1 phase. Centromeres were visualized by CREST sera, and DNA was stained by DAPI. White scale bars indicate 10 µm. (**C**) Quantification of the centromeric GFP-fluorescence intensities of the indicated expressed M18BP1 variants in G1 phase. Centromeres were detected in the CREST channel using a script for semiautomated quantification. Data shown as mean with SD. (**D**) Representative images of fixed HeLa cells showing GFP fluorescence of indicated stably expressed ^GST^M18BP1^GFP^ variants and immuno-stained α-tubulin in mitosis. Centromeres were visualized by CREST sera, and DNA was stained by DAPI. White scale bars indicate 10 µm. (**E**) Quantification of the centromeric GFP-fluorescence intensities of the indicated expressed M18BP1 variants in mitosis. Centromeres were detected in the CREST channel using a script for semiautomated quantification. Data shown as mean with SD. (**F**) and (**G**) Anti-GFP and anti-Tubulin immunoblots of HeLa cell lysates expressing the indicated ^GST^M18BP1^GFP^ variants.

**Table 1:** X-ray data collection and refinement statistics.

|  |  |
| --- | --- |
| Resolution range (Å) | 45.04-2.42 (2.51-2.42)* |
| Space group | C2 |
| a,b, c (Å) | 143.2 120.28 77.36 |
| $\alpha, \beta, \gamma$ (°) | 90.00 121.28 90.00 |
| Molecules/asymmetric unit | 2 |
| Wavelength | 1.0332 |
| Number of crystals | 2 |
| synchrotron | PETRA |
| date | 08. July 2019 |
| Detector | Pilatus6MF |
| **Dataset statistics:** |  |
| Total reflections | 574151 (43031) |
| Unique reflections | 42292 (3857) |
| Multiplicity | 13.6 (11.1) |
| Completeness (%) | 98.97 (90.67) |
| Mean $I/\sigma I$ | 10.37 (1.10) |
| CC1/2 (%) | 99.7 (47.1) |
| R-merge (%) | 18.49 (207.7) |
| R-meas (%) | 19.21 (217.7) |
| R-pim (%) | 5.18 (63.61) |
| Wilson B-factor | 56.74 |
| **Refinement:** |  |
| Reflections in refinement | 42265 (3857) |
| $R_{\text{work}} / R_{\text{free}}$ (%) | 22.91/26.19 |
| Protein atoms | 7320 |
| Ligand atoms | 187 |
| Solvent | 94 |
| average B protein/ligand/solvent (Å<sup>2</sup>) | 64.19/64.63/50.93 |
| Clashscore | 2.48 |
| Rotamer outliers (%) | 2.0 |
| **R.m.s. deviations:** |  |
| Bond lengths (Å) | 0.007 |
| Bond angles (°) | 0.99 |
| **Ramachandran:** |  |
| Favored/allowed regions (%) | 97.68/2.32 |
| Outliers (%) | 0.0 |

\* Statistics for highest resolution shell are shown in parentheses.

